# Genomic exploration of the complex journey of *Plasmodium vivax* in Latin America

**DOI:** 10.1101/2024.05.08.592893

**Authors:** M. J. M. Lefebvre, F. Degrugillier, C. Arnathau, G. A. Fontecha, O. Noya, S. Houzé, C. Severini, B. Pradines, A. Berry, J-F. Trape, F. E. Sáenz, F. Prugnolle, M. C. Fontaine, V. Rougeron

## Abstract

*Plasmodium vivax*, the predominant malaria parasite in Latin America, has a rich and complex colonization history in the region, with debated hypotheses about its origin. Our study employed cutting-edge population genomic techniques, to collect whole genome sequencing data from 620 *P. vivax* isolates, including 107 newly sequenced samples, thus representing nearly all potential source populations worldwide. Analyses of the genetic structure, diversity, ancestry, and also, coalescent-based inferences and scenario testing using Approximate Bayesian Computation, have revealed a more complex evolutionary history than previously envisioned. Indeed, according to our analysis, the current American *P. vivax* populations predominantly stemmed from a now-extinct European lineage, with the potential contribution also from unsampled populations, most likely of West African origin, during post-colonial human migration waves in the late 19^th^-century. This study provides a fresh perspective on *P. vivax* intricate evolutionary journey and brings insights into the possible contribution of West African *P. vivax* populations to the colonization history of Latin America.

## Introduction

*Plasmodium vivax* is the dominant human malaria parasite in most inter-tropical countries outside sub-Saharan Africa, mainly Asia, Middle East, Latin America, and marginally certain regions of Africa [1]. One of the main reasons for its successful geographic expansion is its ability to form dormant stages in human liver cells [2]. This allows *P. vivax* to maintain itself in temperate climates, hiding in a dormant stage during the cold months when the vectors (*Anopheles* mosquitoes) are in diapause, and creating a persistent presence of parasite reservoirs [3]. In Latin America, *P. vivax* is well established from Mexico to Brazil, where it causes 70% of malaria cases [4]. Venezuela, Brazil, and Colombia are particularly affected (79% of human malaria cases in the continent) [4]. Although *P. vivax* is well established in South and Central America and is a major public health issue, the debate is still open on how, when, and from where it colonized this continent. This knowledge is necessary to develop efficient strategies for malaria control and elimination, and for monitoring its spread into new regions.

In recent decades, various methodologies have been employed to trace *P. vivax* history in Latin America with different results. The study of Gerszten *et al.* [5] reported the first detection of *P. vivax* antibodies in the liver and spleen of South American mummies, dating back 3,000 to 600 years before present (ybp), through visualization of species-specific antigens by immunohistochemistry [5]. They suggested that the parasite was already present in Latin America before the arrival of European settlers and the transatlantic slave trade (spanning from 1500 to 1830). However, this result remains debatable. Indeed, antibody cross-reactivity with *Plasmodium* species has been reported [6–8], but this was not tested by the authors. Although molecular techniques are essential to validate this hypothesis [9], these findings raised the crucial question about the route(s) taken by *P. vivax* to colonize Latin America during pre-Columbian period. It would have been unlikely that the parasite followed its human host across the Bering land bridge from Asia to Alaska given the absence of the mosquito vectors necessary for the completion of the parasite’s life cycle at these high latitudes [10]. Nonetheless, the unique ability of *P. vivax* to enter in a dormant state suggests an alternative scenario in which the parasite could have endured pre-Columbian transoceanic voyages from the Asian mainland or western Pacific islands to South America [11]. This hypothesis recently has gained support with the study by Rodrigues *et al.* [12] which documented genetic contributions from Melanesia (a group of islands in Oceania) in the mitochondrial gene pool of *P. vivax* American populations. Despite this evidence, studies using mitochondrial [12] or microsatellite markers [13] consistently indicated African and South-Asian populations as the main genetic sources of the present-day *P. vivax* lineages in Latin America. These studies also suggest the possibility of several independent *P. vivax* introductions into this continent during the transatlantic trade period [12,13].

More recently, the analysis of one *P. vivax* strain extracted from microscope slides from Spain, dating from 1942-1944, gathered as a composite sample called “Ebro”, reignited the debate [14,15]. Indeed, this isolate, due to its genetic proximity to American populations, its position branching at the root of the American clade in the phylogenetic tree, and its estimated divergence coinciding with the transatlantic slave trade period led van Dorp *et al.* [15] to propose that *P. vivax* American populations originated from European colonization during the 15^th^ century (*i.e.* in the post-Columbian period). This study introduced a compelling hypothesis of the European origin of *P. vivax* in Latin America, however, some limitations needed to be considered. First, the study did not consider all possible sources of American *P. vivax* populations. Indee, the dataset included a limited number of *P. vivax* individuals from Africa (n=4), all from Madagascar. As more than seven million slaves, mainly from West/Central Africa, arrived in Central and South America during the transatlantic trade [16], it was imperative to consider West and Central African *P. vivax* populations to precisely determine the colonization history of *P. vivax* in Latin America. The relevance of an African origin must be taken into account because it has now been established that the colonization of Central and South America by *Plasmodium falciparum*, the most virulent human malaria agent, resulted from two independent introductions from West/Central Africa during the transatlantic slave trade [17–19]. The high frequencies of Duffy-negativity in African populations, expected to protect against *P. vivax* infection might exclude such a hypothesis; however, the mounting evidence that *P. vivax* is present in West Africa [20–22], as observed in Mauritania [23,24] and Senegal [25], reopens the door to this primary source. In addition, van Dorp *et al.* [15] only considered four populations from Latin America, thus including a limited number of populations and geographic areas within the parasite distribution range on the continent. Lastly, Dorp *et al.* [15] did not examine the possibility that Ebro represents an introduction of *P. vivax* lineage from the Americas into Europe. This hypothesis was already raised by Escalante *et al.* [26], and hundreds of Europeans migrated to Latin America and then returned to Europe during the 19^th^ and 20^th^ centuries [27].

Therefore, *P. vivax* evolutionary origin in Latin America remains unclear and debated. The objective of this study was to perform a comprehensive population genomic analysis of a novel dataset of *P. vivax* genomes from multiple countries in Latin America and Africa. This study considered a large dataset included 1,133 isolates among which 214 were newly sequenced for this study. After quality filtering, the final dataset included 620 unrelated *P. vivax* genomes from 36 countries worldwide (Fig. 1) among which, there were 107 newly sequenced genomes originating from six countries Latin American countries (Brazil, Ecuador, French Guiana, Guyana, Honduras, and Venezuela), five African countries (Central Africa, Comoros, Ethiopia, Mauritania, and Sudan), three Middle Eastern countries (Afghanistan, Armenia, and Azerbaijan), two South Asian countries (India, Pakistan), and one Oceanian country (Papua New Guinea). The combination of these new genomes with those already available provided a comprehensive assessment of the genomic diversity from a whole nuclear genome perspective by encompassing the present-day *P. vivax* distribution. To our knowledge, this study marks a new and original attempt to address the question of *P. vivax* colonization of Latin America using genetic material from all potential source populations identified in the literature (Europe, Africa, Asia and Oceania). This dataset was used to study the population genetics structure and diversity, the population demographic histories, and to test different evolutionary scenarios of *P. vivax* colonization of Latin America using approximated Bayesian computations. Our results indicated a complex *P. vivax* colonization history in Latin America with multiple waves of migration. Our findings corroborate the European origin proposed earlier [15], but with a more recent colonization date estimated during the post-colonial migration in the 19^th^ century. However, they also emphasize the contribution of an additional unsampled population, that might display a genetic affinity with West African *P. vivax* populations.

**Figure 1.**
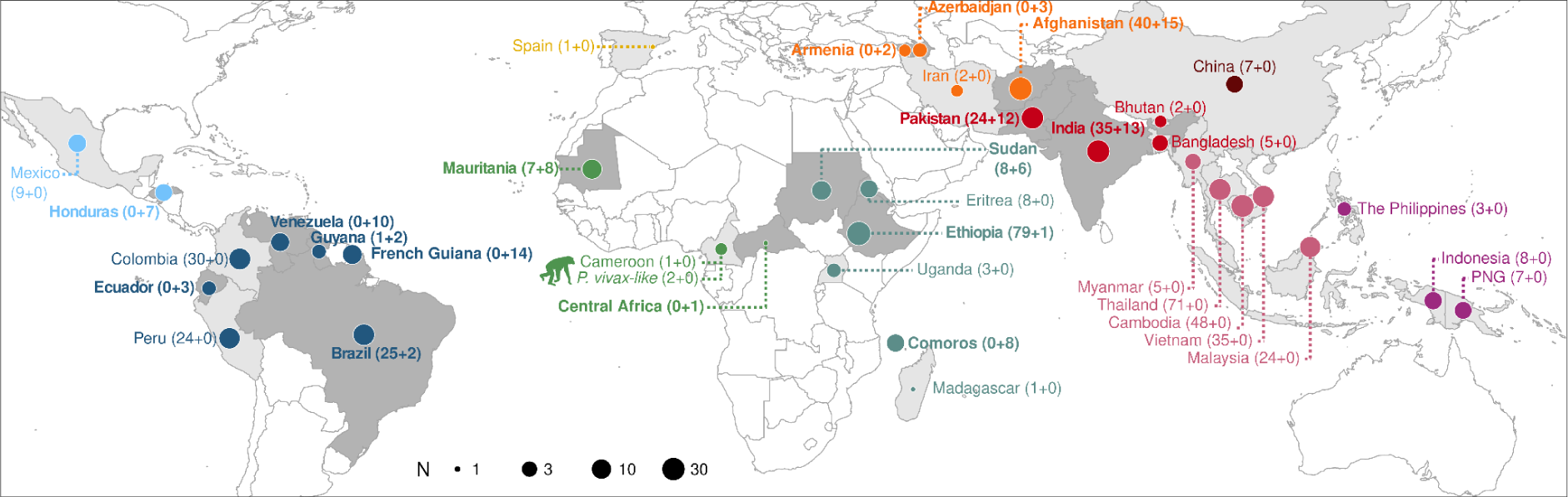
Geographic origin of the 620 human *P. vivax* isolates and two African great apes *P. vivax-like* isolates. Distribution of the samples studied per region: Central America (n = 16, light blue), South America (n = 111, dark blue), Europe (n=1), West Africa (n = 18, light green), East Africa (n = 115, dark green), Middle East (n = 62, orange), South Asia (n = 91, light red), East Asia (n = 7, dark red), Southeast Asia (n = 183, pink), and Oceania (n = 18, purple). The circle size is proportional to the sample size in log_10_ units. The countries included in the sampling effort are differentiated by varying shades of gray. : darker gray for countries where new samples were sequenced and lighter gray for countries where already published samples were used. Country names highlighted in bold indicate that new genomes were sequenced in this study. In brackets, the number of samples from the literature is listed first, followed by the number of newly sequenced samples. The chimpanzee pictogram indicates the two great apes *P. vivax-like* samples studied. PNG: Papua New Guinea.

## Results

### A worldwide genomic sampling of *P. vivax*

In this study, 214 new *P. vivax* strains from Central and South America, West, Central and East Africa, Middle East, South Asia, and Oceania were sequenced (for more details see Materials and Methods; S1 Table). Then, these new genomes were combined with 919 modern genomes from the literature [28–30] (see Materials and Methods for more details; S1 Table, S1 Fig.) and one ancient sample (Ebro) from Spain sequenced by van Dorp *et al.* [15]. Among the 1,133 modern samples across the globe, genomes with >50% missing data (n=68) and those with multi-strain infections (*i.e.*, multiple *P. vivax* genotypes), reported with a within-sample infection complexity (*F_WS_*) >0.95 were removed (n=320; S2 Fig.). In addition, for all strain pairs in each country displaying excessive relatedness (pairwise-identity by descent, IBD >0.5), one strain for each pair was removed (n=124; S2 Fig.). The final dataset included whole genome sequencing data of 619 modern *P. vivax* samples from 34 countries, one ancient DNA sample (Ebro) from Spain (Fig. 1 and S1 Table) and two genomes of *P. vivax-like* from Cameroon, isolated from chimpanzees [29] and used as outgroup. The mean sequencing depth ranged from 2.88 to 1,484.71 X for the modern genomes and the mean sequencing depth was 1.13 X for Ebro (S2 Table). The details about the bioinformatic pipelines, data processing, and the number of single nucleotide polymorphisms (SNPs) used are described in Materials and Methods and in S1 and S3 Figures. Some of our analyses considered the genotype likelihood using the ANGSD software ecosystem [31], rather than the classic SNP calling followed by hard quality filtering, to account for the genotyping uncertainty related to the sequencing depth heterogeneity. This strategy using ANGSD allowed considering 20,844,131 sites across the 620 genomes analyzed. Nevertheless, as some analyses required actual SNP calling followed by appropriate quality filtering, a Variant Call Format (VCF) dataset composed of 950,034 SNPs was also generated (for more details refer to Materials and Methods and S1 and S3 Fig.).

### Genetic relationships of the American *P. vivax* with worldwide populations

To explore the genetic relationships of Latin American *P. vivax* isolates with worldwide *P. vivax* strains, the population structure patterns of 620 *P. vivax* isolates collected in 36 countries (Fig. 1) were investigated, using complementary population genomic approaches (see the Materials and Methods and S1 Fig.): (i) a principal component analysis (PCA) using *PCAngsd* [32], (ii) a model-based individual ancestry analysis using *PCAngsd* [32], and (iii) a maximum likelihood (ML) phylogenetic tree using *IQ-TREE* [33]. All three approaches showed consistent results splitting first the genetic variation geographically into four distinct genetic clusters (Fig. 2): (1) Oceania, East and Southeast Asia, (2) Africa, (3) Middle East and South Asia, and (4) Latin America.

**Figure 2:**
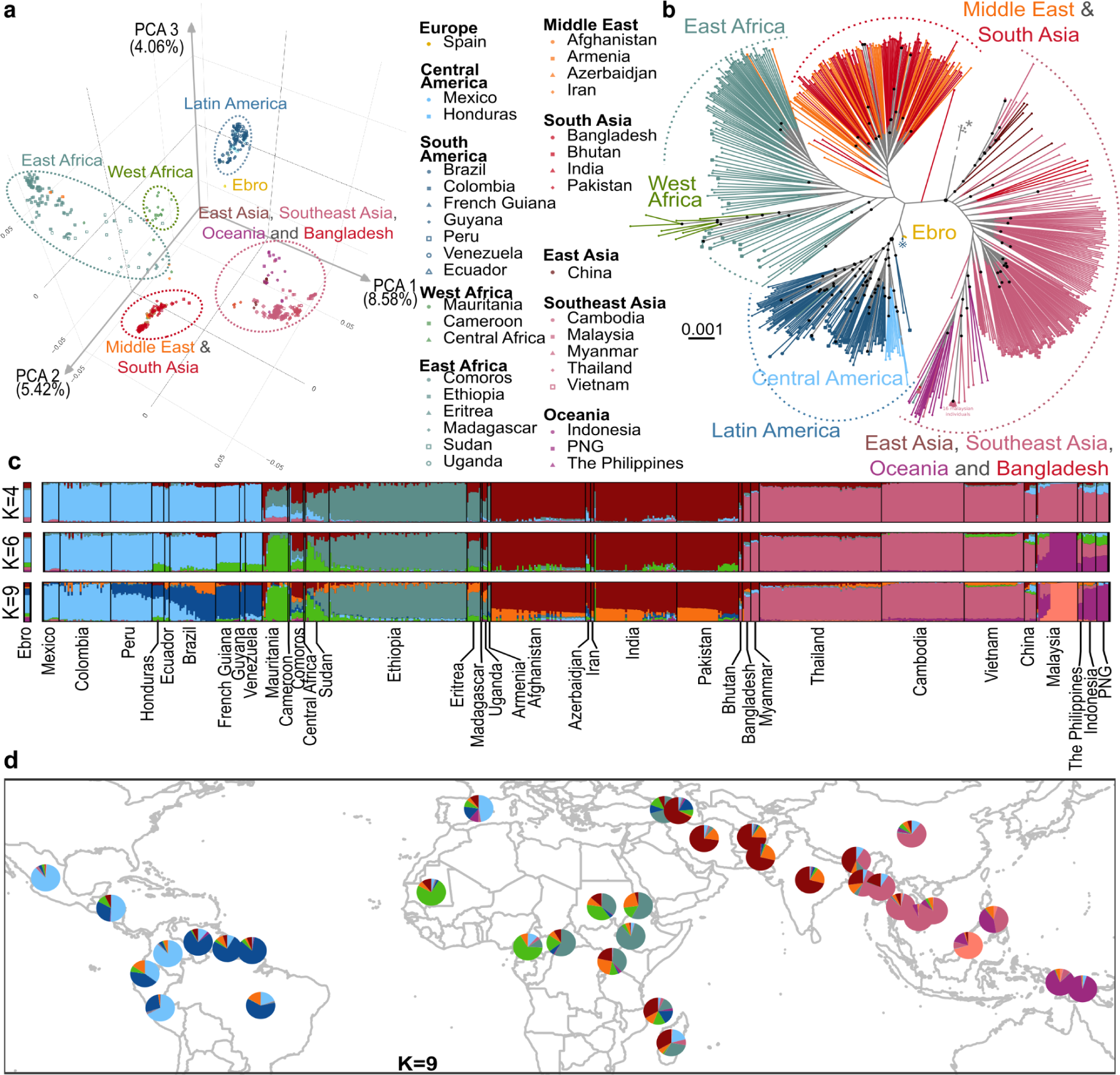
Worldwide genetic structure of *P. vivax*. **(a)** Principal component analysis of 619 modern *P. vivax* strains and the Ebro ancient DNA sample from Spain, showing the first, second, and third PCs based on the genotype likelihood of 105,527 unlinked SNPs (see S2 Fig. for more details). **(b)** Maximum likelihood (ML) tree of the 620 *P. vivax* individual genomes obtained with IQ-TREE [33] based on a general time reversible model of nucleotide evolution [34], as determined by ModelFinder [35]. The ML tree includes two *P. vivax-like* strains from African great apes indicated by an asterisk (*) used to locate the root. Note that the length of the outgroup branch was truncated. Black dots at nodes correspond to highly supported nodes (SH-aLRT ≥ 80% and UFboot ≥ 95%). The reference mark (※) highlight a Brazilian *P. vivax* isolate, discussed in the text. **(c)** Individual genetic ancestry assuming K=4 and K=9 distinct genetic clusters estimated using *PCAngsd* (see S5 Fig. for the other K values). **(d)** Population mean genetic ancestry at K=9 estimated using *PCAngsd* showing the same ancestral proportions. PNG: Papua New Guinea.

In the PCA, the first three principal components (PC) revealed these four major genetic clusters (Fig. 2a and S4 Fig.). PC1 separated Oceania, East and Southeast Asia from Africa, the Middle East, South Asia, and Latin America. PC2 splitted American populations from all the others. African populations split from all the other populations on PC3. The same genetic structure with four genetic clusters was also given by the ML tree (Fig. 2b). *P. vivax* populations from East and Southeast Asia, Bangladesh, and Oceania split first and cluster closer to the outgroup and apart from those from the rest of the world. Populations from Latin America, Middle East-South Asia and Africa clustered in three distinct lineages, an the last two were more closely related to each other than to American lineages. Patterns of individual genetic ancestry (Fig. 2c and S5 Fig.) were also in line with the PCA and ML tree results, displaying again four distinct genetic clusters. As the number of genetic clusters tested (K) increased, the four different genetic clusters also became distinct in terms of genetic ancestry. All the populations from East and Southeast Asia, Bangladesh and Oceania split first from the others at K=2. Then, populations from Latin America split from from Middle East-South Asia and from Africa, which became differentiated only at K=4 (Fig. 2c).

Data analysis at a finer sub-structuration level at K=6 (the best K value based on the broken-stick eigenvalues plus one, following the recommendations of Meisner and Albrechsten [32]) (Fig. 2c and S5 Fig.) showed that Latin America remained as a single genetic cluster well separated from the rest of the world. Asia displayed three clusters (South Asia-Middle East, East Asia-Southeast Asia, and Oceania-Malaysia) and Africa was divided into East and West Africa. Noteworthy, several American populations (Honduras, Ecuador, Brazil, Venezuela, French Guiana, and Guyana) shared genetic ancestry with the cluster that contains West African isolates (in green, Fig. 2c).

Only at K=9, an additional sub-structuration of the genetic clusters was observed among the American *P. vivax* populations (Fig. 2c). Specially, two distinct genetic clusters were identified, as previously observed [13,29,36]: (1) isolates from Mexico, Honduras, and Colombia gathered together as a Central American group, and (2) isolates from the French Guiana, Guyana, and Venezuela formed another distinct cluster as an Amazonian group (Fig. 2d). The other populations from Brazil, Peru and Ecuador displayed patterns of admixed genetic ancestry between these genetic clusters.

In Africa, *P. vivax* populations were distributed in two distinct genetic clusters, one composed of the Mauritanian and Cameroonian samples, and the other including samples from Ethiopia and Eritrea. Sudan and Central African samples displayed admixed genetic ancestry between these African clusters (Fig. 2d).

In Asia, a subset of the Malaysian population (n=16) exhibited no evidence of admixture with other populations. These Malaysian samples were closely clustered in the ML tree (Fig. 2b) and on PC 4 were separated from the other Asian populations (S4 Fig.). This pattern has already been reported before in this region [37] and it could be due to a sharp decline in *P. vivax* prevalence in Malaysia over the past decade [38,39], that might have increased the population genetic drift.

In the PCA, the ancient DNA sample from Spain (Ebro) appeared to be genetically closer to the American populations than to any other population (Fig. 2a), with which it shared a similar ancestry pattern (Fig. 2c-d). In the ML tree, it clustered with one Brazilian sample (indicated with this symbol ※ in Fig. 2b), the oldest sample from the Latin America dataset, collected in 1980 [30]. Similarly, in a previous study based on mitochondrial DNA, it clustered with South American samples [14]. Nevertheless, the nodes clustering these individuals were poorly supported in the ML tree (Fig. 2b), preventing any strong interpretation of the origin of Ebro.

Overall, the population structure analyses (PCA, ancestry plots, and ML phylogenetic tree) supported the hypothesis that the Latin American *P. vivax* populations are genetically closer to the ancient DNA Ebro isolate from Spain, followed by the present-day populations from Africa and from Middle East & South Asia. Noteworthy, both the American and Spanish Ebro genomes displayed some admixed genetic ancestry with other populations from the rest of the world. For example, at K=6 and 9 in the Admixture analysis (Fig. 2c and 2d), Ebro shared approximately 50% of genetic ancestry with the American populations (light blue), but the rest originated from populations located in West Africa and Middle East (respectively light green and brown). Some American populations (Honduras, Ecuador, Brazil, Venezuela, French Guiana, and Guyana) also displayed shared genetic ancestry, particularly with isolates from West Africa.

### Admixture and population branching order between American *P. vivax* and the rest of the world

To further investigate potential admixture events and identify candidate source populations that could have contributed to *P. vivax* colonization of Latin America, first computed population admixture graphs were computed using two complementary approaches implemented in *TreeMix* [40] and in *AdmixtureBayes* [41].

The *TreeMix* analysis determined that the optimal number of migration edges was two or five (S6 Fig.), while *AdmixtureBayes* analyses converged toward an optimal solution with two admixture events (S7 Fig.). Concerning the genetic structure analyses (Fig. 2), both methods identified the four previously described genetic clusters (Fig. 3): (1) Oceania, East and Southeast Asia, (2) Africa, (3) Middle East and South Asia, and (4) Latin America.

**Figure 3:**
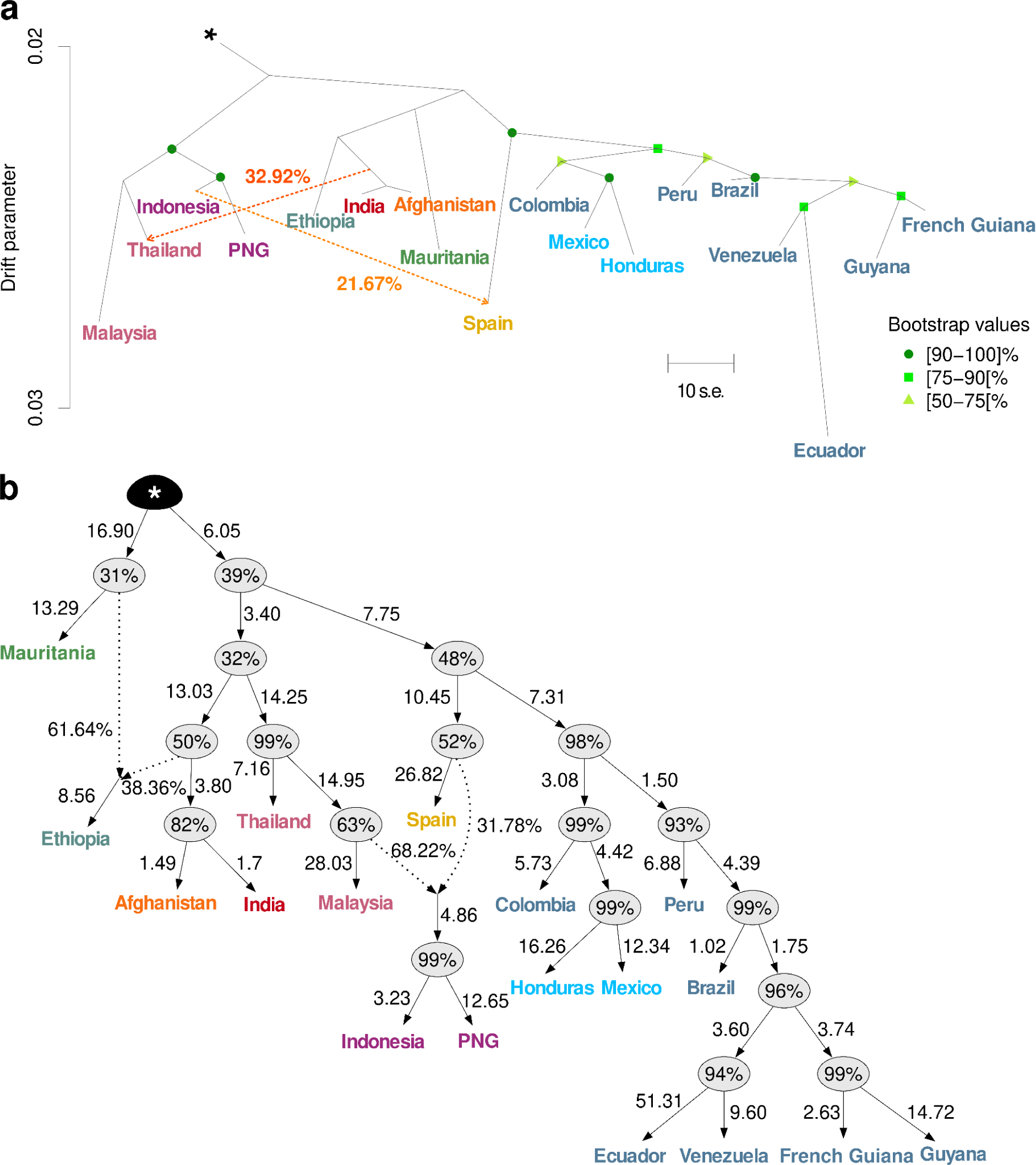
Population graphs describing the genetic relationships and admixture proportions between *P. vivax* populations. **(a)** Population tree estimated using *TreeMix* for a subset of 18 *P. vivax* populations with two migration edges (orange arrows) and rooted using the two *P. vivax-like* genomes indicated with the asterisk (*). The scale bar shows ten times the mean standard error (s.e.) of the covariance matrix. The migration weight is indicated as a percentage in orange on the arrows. **(b)** Network topology of a subset of 18 *P. vivax* populations with the highest posterior probability obtained with *AdmixtureBayes*, rooted with teh two *P. vivax-like* genomes (*). The branch length indicates the genetic divergence between populations (measured by drift), multiplied by 100. The percentages in the nodes are the posterior probability that the true graph has a node with the same descendants. For each admixture event (indicated by the dotted arrows) the percentages illustrate the admixture proportion. PNG: Papua New Guinea.

The American populations grouped into a single highly supported monophyletic cluster (≥75% bootstrap support in *TreeMix*, Fig. 3a, and ≥98% posterior probability in *AdmixtureBayes*, Fig. 3b). This suggests that American populations most likely originated from a single introduction or multiple introductions from genetically similar, but possibly admixed source populations from the rest of world. This second hypothesis is supported by the poorly resolved branching pattern close to the root in the *TreeMix* and *AdmixtureBayes* analyses (Fig. 3). The branching patterns obtained in both population graphs showed a finer substructuration of the American populations (Fig. 3): one cluster grouping Mexico, Colombia, and Honduras populations, another one with the French Guiana, Guyana, Venezuela and Ecuador populations, and in between, Brazil and Peru populations branched at an intermediate position reflecting their admixed genetic ancestry observed in the genetic structure analyses (Fig. 2).

The ancient Spanish sample (Ebro) stood out as the isolate most closely related, but genetically distinct, from the American cluster, branching at its base. This ancient isolate had a notably long branch that reflected important genetic drift compared with other present-day populations. Such a long branch for Ebro may be an artifact because it is a single haploid sample from an ancient population [42]. As rare and common alleles in that ancient population could not be properly estimated, this could have led to an inflated estimate of the genetic drift as previously reported [42]. Additionally, the *TreeMix* and *AdmixtureBayes* analyses detected an admixture event between Ebro and Oceanian populations, corroborating the shared genetic ancestry identified in the population genetic structure analyses (Fig. 2d-c). The *TreeMix* analysis also identified a migration edge from the common ancestor of India and Afghanistan into Thailand.

To assess potential admixture events between the American populations and those from the rest of the world, the admixture *f_3_*-statistics were computed. This involved testing American populations against potential source populations from both within Latin America and from all other geographical regions worldwide. None of the tested *f_3_* combinations yielded a significantly negative result, as expected if admixtures occured between the American populations and one of the two tested sources (S8 Fig.). As underlined by the authors of the *f_3_* admixture test [43], these non-significant results do not preclude that admixture could have occurred in a distant past, with a residual signal remaining only in a minor fraction of the genome. This would be also expected if the source population(s) that founded the Latin American populations were already admixed.

### Effective population size changes and split time of American *P. vivax* populations

The next step was to determine whether colonization of Latin America by *P. vivax* was associated with founder effects and population bottlenecks, as can be expected when colonizing a new environment. To this aim, the nucleotide diversity (π) as well as the Tajima’s D values [44] where computed in the core genome, considering only populations with at least 10 samples. The π values in the American populations (S9 Fig.) were marginally lower compared with those observed in other populations, with exceptions in Malaysia and Ethiopia. In Latin America, the π values displayed distinct patterns: populations along the Pacific coast (Colombia and Peru) exhibited significantly lower median π values than population in the Amazon (Brazil, French Guiana, and Venezuela)(∼8×10^-4^ versus ∼1×10^-3^, *p-value* <0.001, Wilcoxon signed-rank test and Bonferroni correction, S9 Fig.). The Tajima’s D distributions in the same *P. vivax* populations were largely negative, indicating excess of rare variants over shared variants, consistent with demographic expansions. However, the Tajima’s D distribution of populations from Colombia, Peru, and Ethiopia were closer to zero, with a notable presence of positive values (S9 Fig.). Overall, these descriptive statistics are consistent with the demographic expansion of the American populations. They also suggest that no strong bottleneck occured during *P. vivax* invasion history of Latin America or that the founding isolates were somewhat admixed and harbored genetic diversity comparable to that of the source populations.

The changes in effective population size (*N_e_*) with time were inferred using the coalescence rates (CR) estimated with the software *Relate* [45] for modern samples. Consistently with the π and Tajima’s D values, these analyses showed that all modern populations across the world displayed similar trends, characterized by a monotonic decline in *Ne* values that started roughly ∼100,000 generations ago (∼20,000 years ago, assuming a generation time of 5.5 generations/year [29] and a mutation rate of 6.43 x 10^-9^ mutations/site/generation [29]) followed by a recent and moderate expansion (Fig. 4a). Noteworthy, all American *P. vivax* populations displayed slightly lower *Ne* values than the other populations, and only three populations from Africa and Oceania showed comparable trends (Mauritania, Malaysia, and Papua New Guinea). The *Ne* values of all American populations became noticeably distinct from the other populations since the last ∼500 to 2,000 generations (∼100-300 years ago). This time range suggests a putative divergence time of the American populations from the other populations, but also with each other. Besides the trees (Fig. 2b and 3) where the American populations formed a single well-supported cluster, the synchronous departures of the *Ne* values of the American populations, compared with those from other populations worldwide, also suggests a single introduction from one source or multiple introductions from genetically similar sources, possibly admixed, in Latin America.

**Figure 4:**
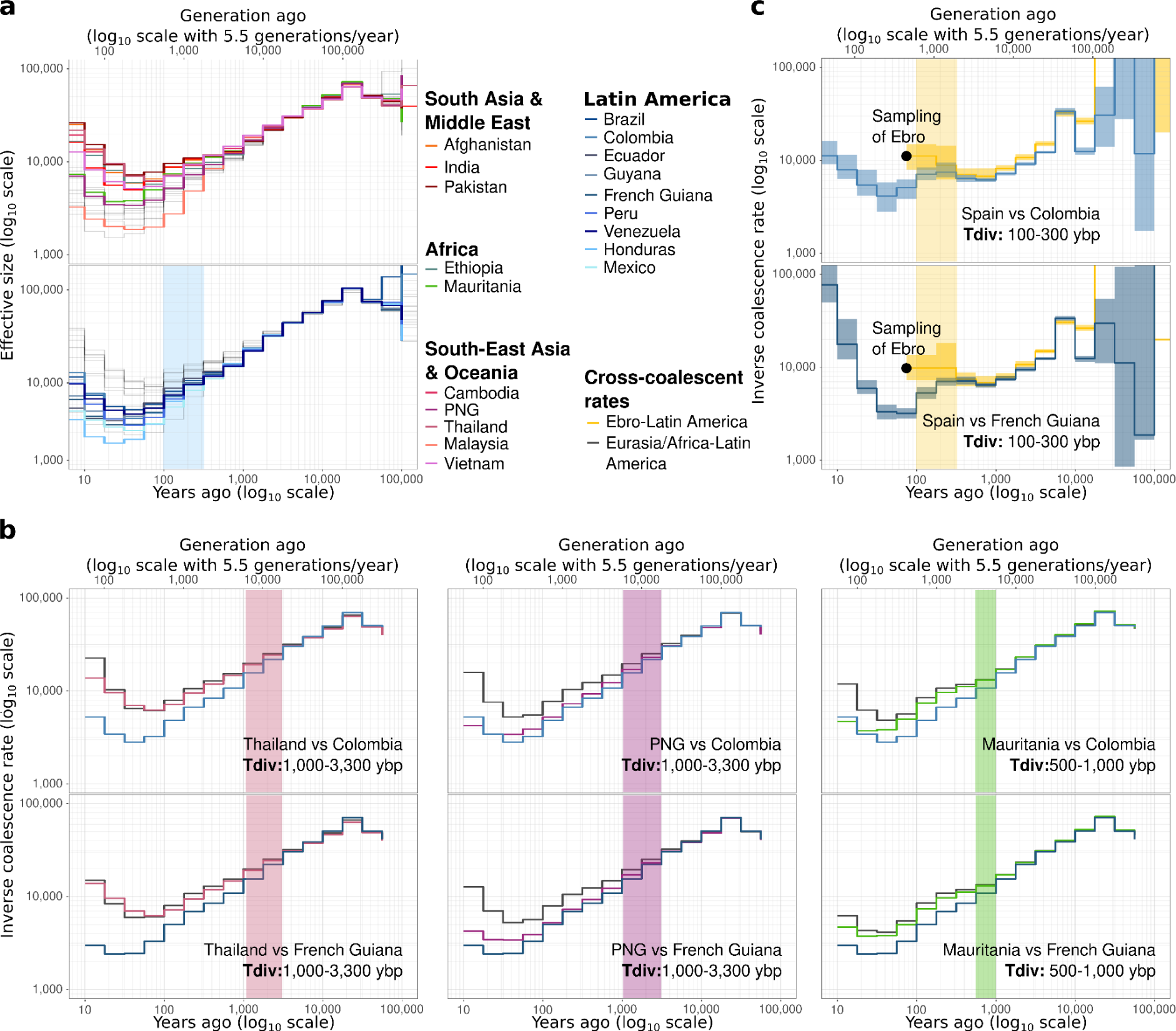
Coalescent-based inference of the demographic history of *P. vivax* populations. **(a)** The variation of the effective population (N_e_) size was estimated using *Relate* (axes are log_10_ transformed). The period highlighted by the blue rectangle corresponds to the diversification of the Latin American populations. **(b)** Comparison the of inverse coalescence rates and inverse cross-coalescence rates from *Relate* between Latin America (Colombia on top, French Guiana below) and Eurasia/Africa (left to right: Thailand, Papua-New-Guinea, Mauritania). The axes are log_10_ transformed. The periods highlighted by the colored rectangles correspond to the divergence time (*Tdiv*) between populations and are also specified in the panels. **(c)** Comparison of the inverse coalescence rates and inverse cross-coalescence rates between Latin America (Colombia on top, French Guiana below) and Ebro. The smoothing line shows the 95% confidence interval resulting from 100 bootstraps, inferred using *Colate*. The dot indicated Ebro sampling time. The axes are in the log_10_ scale. The period highlighted by the light-yellow rectangles corresponds to the divergence time (*Tdiv*) between the American populations and Ebro and is specified. PNG: Papua New Guinea.

The divergence time estimates were refined using the inverse cross-coalescent rate (CCR) between representative populations from the American clusters (Colombia and French Guiana) and other populations in the world that are representative of distinct genetic clusters (Fig. 2 and 3), as well as the ancient isolate from the Ebro region in Spain. This analysis used the program *Relate* [45] for modern samples (Fig. 4b) and the program *Colate* [46] for the ancient Ebro sample (Fig. 4c). The split time between populations can be estimated by comparing the changes in CCR and CR values between populations and within population respectively. The split time corresponds to the time when the inverse of the CR values within each population departs from each other and the inverse of the CCR values between the populations increases. This means that the CR between populations decreases, and therefore they are diverging [47]. According to the CCR variation (Fig. 4b), the divergence times estimated between the American populations and Asian (*i.e.* Thailand) or Oceanian (*i.e.* PNG) populations occurred more than 1,000 ybp (or more than 55,000 generations ago). The divergence time between populations from West Africa and Latin America dated back 500-1,000 ybp (∼3,000-5,500 generations ago). Lastly, the split between the American populations and the ancient Ebro isolate from Spain would have been more recent (100-300 ybp or 550-1,650 generations ago). These divergence time estimates are consistent with the population branching order from the previous analyses (Fig. 2b and 3).

### The role of West African and European *P. vivax* populations in the invasion of Latin America

Our analyses of the population structure and relationships between populations indicated that American *P. vivax* populations form a highly supported monophyletic cluster (Fig. 2b and Fig. 3). These results and the concomitant divergence of all American populations (Fig. 4a) suggested that they descend from a single introduction from one source or from multiple introductions from genetically similar and possibly admixed sources. The ancient Ebro sample from Spain was the isolate the most closely related to the American populations. It displayed an admixed genetic ancestry, half of which was shared with the American *P. vivax* populations, and the other half originated from multiple sources (Africa, the Middle East, and even Asia) (Fig. 2c). Nevertheless, West African *P. vivax* populations also may have contributed to the genetic make-up of the Latin American populations. This is suggested by the recent human history that involved massive population movements between Europe, West Africa and Latin America [16]. This is also supported by the genetic ancestry analyses showing that to some extent, West African populations also may have also contributed to the genetic ancestry of *P. vivax* populations in Latin America (Fig. 2c and S5 Fig.). However, no evidence of admixture was detected in the population graph analyses between American and African populations (Fig. 3, S6 and S8 Figures). Furthermore, American populations did not share ancestry with Asian or with Oceanian populations (Fig. 2c), and the divergence times with Asian and Oceanian populations were much older than with West African or European population (Fig. 4). The European ancient Ebro isolate and the West African populations were the only groups from Eurasia and Africa that showed credible evidence of shared genetic ancestry with the American populations. Therefore, it was important to formally test their putative roles as source populations in the *P. vivax* invasion history of Latin America.

We used the simulation-based approximate Bayesian computation (ABC) statistical framework that relies on the Random Forest tree classifier (ABC-RF), as implemented in DIYABC-RF [48]. This allowed comparing distinct scenarios of invasion history of Latin America that differed in terms of population branching order, admixture events, time of split and admixture, and that involved or not unsampled (or ghost) populations (see Fig. 5a and Material and Method for details). In an ABC-RF analysis, genetic data are simulated in different demographic scenarios (see parameters in S3 Table), and summary statistics from the resulting simulated data are compared with those obtained from the observed data. This analysis considered twelve colonization scenarios that involved three representative populations: one from West Africa (Mauritania), one from Europe (the Ebro sample), and one from America (Colombia). An unsampled (ghost) population was also used in scenarios 5 to 9, 11, and 12 (Fig. 5a). Scenario 2 considered only a European origin; scenarios 3, 4 and 12 considered only an African origin; and scenarios 1, 6, 7, 8, 9, 10 and 11 considered some admixed histories.

**Figure 5:**
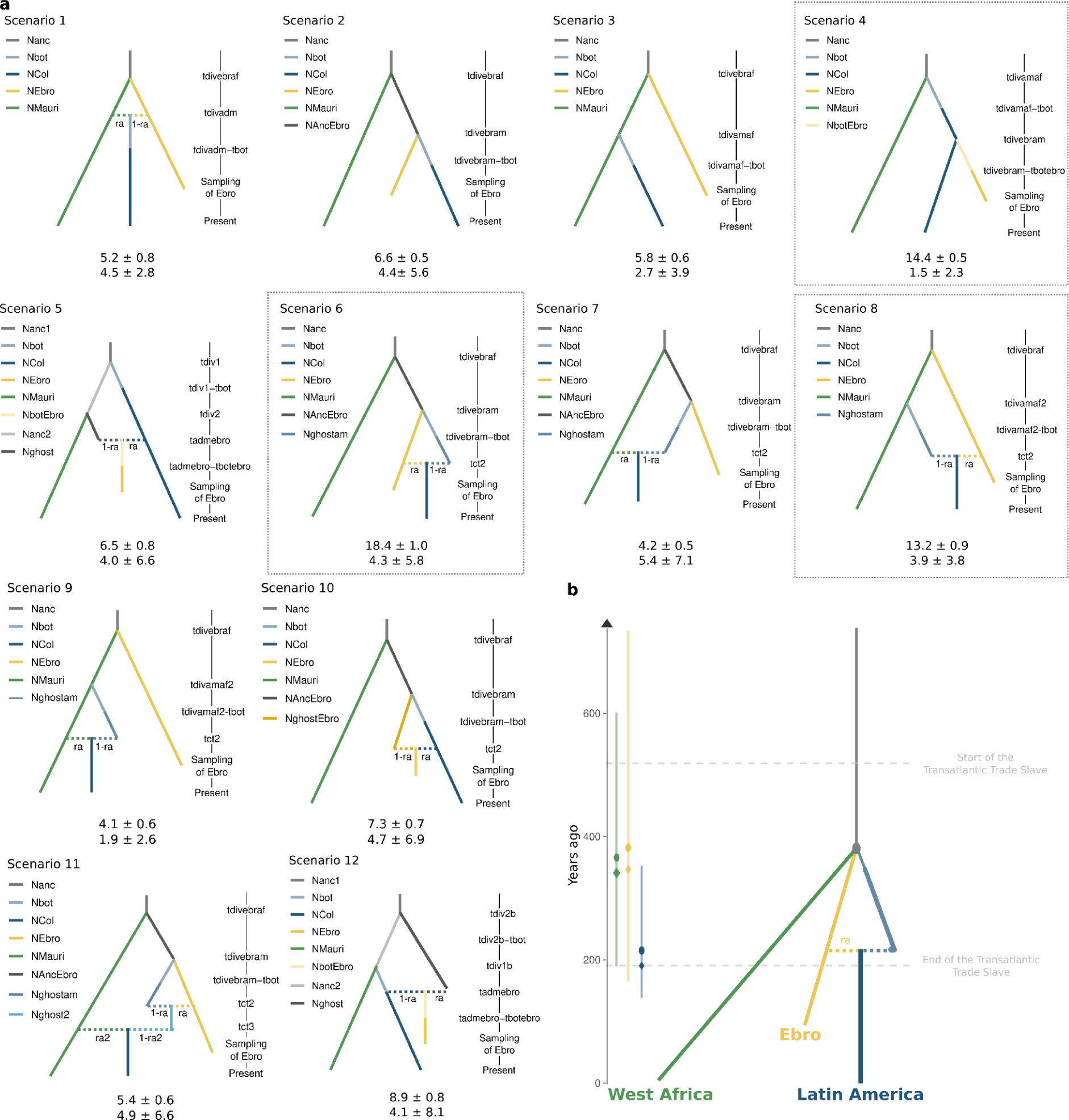
Most likely *P. vivax* colonization history scenarios of Latin America inferred using the Approximate Bayesian Computation Random Forest approach. **(a)** Twelve possible colonization scenarios. Solid line colors correspond to population size parameters and horizontal dashed lines represent admixture events. Time parameters are on the right (not to scale). For the distribution of each parameter, refer to the S3 Table. Under each scenario is the mean percentage of votes ± standard deviation (SD) and the mean type II error ± SD. The three most supported scenarios (scenarios 6, 4 and 8, with 46.0% of the total votes) are highlighted by dotted-line rectangles. **(b)** The best-supported scenario (scenario 6) in the DIYABC-RF analysis, scaled to relative time-parameter estimates (converted to years assuming a generation time of 5.5 generations/year). On the left, each time parameter estimate is indicated by the mean (circle), median (diamond), and 90% confidence intervals (colored bars).

The most likely scenario identified was scenario 6 (Fig. 5) in which the American *P. vivax* populations originated from an admixture event between the European and an unsampled population derived from an ancestor to the European gene pool that branched with West African ancestral populations. This scenario was the most strongly supported with a mean posterior probability (± standard deviation) of 46.2 ± 1.3% and a mean percentage of RF classification votes of 18.4 ± 1.0% votes in the ABC-RF analysis (Fig. 5a). Scenarios 4 and 8 were the two other best scenarios (mean percentage of RF classification votes of 14.4 ± 0.5% and 13.2 ± 0.9%, respectively). In scenario 4, Ebro could be a migrant from Latin America, while the American populations would descend from an ancestral population of West African origin. In scenario 8, the American populations would have derived from an admixture event between the European populations and an unsampled population of direct West African origin. It displayed a colonization scenario close to scenario 6. Combined, these three scenarios (6, 4 and 8) collected 46.0% of all votes, while each of the other nine scenarios received less than 8.9% of all votes (Fig. 5a). Ten replicates of the ABC-RF analysis supported strongly and unanimously this result. Furthermore, the confusion matrix (S10 Fig.) and the prior error rate of 42.1 ± 0.0% indicated the overall performance of the RF classification analysis and highlighted high power to distinguish among the twelve competing scenarios.

In the optimal scenario 6, model parameter estimates indicated that the divergence time between the ancestral population of Ebro and the West African population (in green in Fig. 5b) was approximately 363 years ago (or 2,017 generations ago). Similarly, the estimated divergence time between the unsampled (ghost) population and Ebro (in yellow in Fig. 5b) was approximately 379 years ago (or 2,108 generations ago). Therefore, the ABC-RF analysis suggested that the two split times between West Africa, Europe and the unsampled population were nearly synchronous. These split time estimates from the ABC-RF are close to those from the CCR estimates between the American populations and those from Europe (Ebro) and West Africa (Fig. 4). Furthermore, the admixture event between the unsampled population and the European population (in yellow in Fig. 5b) most likely occurred 212 years ago (or 1,179 generations ago), with a predominant European contribution (*ra* = 60%).

Altogether, these ABC-RF results support the hypothesis that American *P. vivax* populations descended from now-extinct European populations, but also from unsampled population(s), possibly originating from West Africa, an event that happened during the post-colonization migration of the American continent (second half of the 19th century). This analysis did not fully rule out the American origin of the European Ebro sample, as suggested by scenario 4 (Fig. 5).

## Discussion

The investigation of the colonization of Latin America by *P. vivax* has been approached through a variety of methodologies. Previous studies using mitochondrial DNA [12] and immunohistochemistry [5] suggested the existence of a pre-Columbian presence and potential Asian/Oceanian origins. Other studies using mitochondrial DNA [12] or microsatellites [13] indicated a potential influence from West Africa during the transatlantic slave trade era. Recently, van Dorp *et al.* [15] analyzed complete *P. vivax* nuclear genomes that included a composite sample from a ancient slide, and suggested a European introduction that coincided with the European colonization of Latin America. However, this study had important limitations, particularly (1) the absence of samples from West Africa, the region from where millions of enslaved persons were transported to Latin America during the transatlantic slave trade [16] and the source of American *P. falciparum*, another human malaria parasite [17–19]; (2) the European origin hypothesis relied only on a single, very low coverage, and composite sample originating from a multi-*Plasmodium* species co-infection (Ebro sample). Therefore, the possibility that the Ebro sample were of American origin and transferred back into Europe during the massive human movements between Europe and Latin America in the 19^th^ and 20^th^ centuries cannot be excluded, as suggested by Escalante *et al.* [26].

Considering these factors, our study conducted population genomic analyses were based on a novel dataset of 620 *P. vivax* isolates from 36 countries, including 107 newly sequenced samples, particularly from Latin America and West Africa. To the best of our knowledge, this dataset represents the first comprehensive collection of all potential source populations proposed in the literature, thereby allowing the exploration of the evolutionary history of *P. vivax* colonization of Latin America. In addition to population structure and demographic history analyses, the influence of European and African populations in the colonization of the American continent was tested using ABC scenarios.

### *Plasmodium vivax* genetic structure and diversity in Latin America

The analysis based on PCA, ancestry plots, ML phylogenetic tree, and population admixture graphs revealed a consistent genetic structure among the Latin American *P. vivax* populations within the Americas and as well as to *P. vivax* populations worldwide, as previously described [29,36,49]. On a worldwide scale, four distinct genetic clusters corresponding to different geographical regions were identified: (1) Oceania, East, and Southeast Asia; (2) Africa; (3) Middle East and South Asia; and (4) Latin America (Fig. 2 and 3). As previously reported [13,29,36], in Latin America, two separate genetic clusters were identified, representing the Central American region (Mexico, Honduras, and Colombia) and the Amazonian region (French Guiana, Guyana, and Venezuela). Genetic structure analyses also indicated the presence of an admixed cluster that included *P. vivax* populations from Brazil and Peru (Fig. 2c-d and Fig. 3), thus suggesting potential a gene flow between these clusters in Latin America. These two distinct genetic clusters and the potential gene flow had been previously described. For instance, Sutanto *et al.* [50] established a genetic connection between Colombian and Peruvian isolates, indicating a possible cross-border spread between theses countries. Other studies confirmed genetic links between the various ancestral populations previously identified in Brazilian and Peruvian *P. vivax* populations, suggesting significant gene flow between these countries [36], as well as with other Amazonian countries such as Guyana [36], French Guiana and Venezuela [13]. These results highlight the possibility of gene flows between Peru, Brazil, Guyana, Venezuela, and French Guyana that could be explained by the large asymptomatic human reservoirs and the dormant stage characteristic of *P. vivax* [51–53].

The geographic distribution of genetic diversity across the Latin American region revealed that *P. vivax* populations along the Pacific coast (Colombia and Peru) exhibited lower genetic diversity (π) compared with Amazonian populations (Brazil, French Guiana, and Venezuela) (S9 Fig.). This is in line with previous studies [36,54], and could stem from transmission pattern variations across regions [36]. Indeed, coastal areas experience more periodic malaria transmission events compared with the Amazon region, where transmission is more stable throughout the year [55,56]. However, some studies found that *P. vivax* populations can exhibit high genetic diversity during periods of low transmission rates [54,57]. This means that data on spatial variation in genetic diversity must be interpreted with caution.

### Shared genetic ancestry between *P. vivax* from Latin America, Europe, and West Africa

The population genetic structure analyses (PCA, ancestry plots, and ML phylogenetic tree) showed that *P. vivax* populations from Latin America are genetically closer to the ancient DNA Ebro isolate from Spain, followed by present-day populations from Africa, Middle East and South Asia (Fig. 2). Noteworthy, both the American isolates and the Ebro isolate displayed some admixed genetic ancestry with other populations from the rest of the world. This suggests that European *P. vivax* might not be the only genetic source involved in the colonization of the American continent. When considering the admixture analysis, >50% of the Ebro genetic ancestry was shared with the American populations (light and dark blue), but the rest originated from West African and Middle Eastern population (light green and brown, Fig. 2c-d). In addition, some American populations (Honduras, Ecuador, Brazil, Venezuela, French Guiana, and Guyana) also displayed shared genetic ancestry with those from West Africa. These results support European participation in the genetic make-up of Latin American *P. vivax* populations, but also indicate that the history of the invasion of Latin America could have been more complex with the potential involvement of West African genetic sources.

### No genetic evidence of an Asian contribution to the Latin American *P. vivax* populations

Previous studies suggested that Asian *P. vivax* populations might have contributed to the colonization history of Latin America. For instance, a study based on mitochondrial genomes found evidence of genetic ancestry from Oceanian *P. vivax* strains in American populations [12]. Our genomic analysis also suggested a potential introgression from Indonesia to Mexico, as shown by the *TreeMix* results. However, this introgression appears to be minimal, close to 0% (S6 Fig.), and was not corroborated by the *f_3_*-statistics (S8 Fig.). This introgression could have been the consequence of some historical introductions of Asian *P. vivax* during human migrations, but it could also mirror part of the history of the European *P. vivax* populations, that were the main sources of American *P. vivax*. Indeed, the population graph analysis of *TreeMix* and *AdmixtureBayes* (Fig. 3) detected admixture events that involved Ebro and the Oceanian populations (Papua New Guinea and Indonesia). These admixture events may be due to genetic exchanges between European and local parasites from the Philippines, an archipelago of islands close to Indonesia that was a Spanish colony for more than three centuries (from 1565 to 1899). The reported contribution of Melanesian *P. vivax* isolates to American *P. vivax* genomes may reflect historical introgression, admixture events, or incomplete lineage sorting within the Spanish genomes. Therefore, the likelihood of Asian *P. vivax* serving as genetic sources in the colonization of Latin America appears relatively low, and possibly also integrated within the history of European populations.

### Latin American *P. vivax* populations shared genetic ancestry with the now-extinct European population, but not only

As the Ebro sample showed closer genetic ties with American *P. vivax* populations than with Middle East populations (Fig. 2-3), several scenarios where Ebro came from Latin America (scenarios 4, 5, 10 and 12 in Fig. 5a) and scenarios where Ebro contributed to the genetic make-up of Latin America (scenarios 1, 6, 8 and 11 in Fig. 5a) were tested. The best scenario (scenario 6: 18.4% of votes) suggested that European *P. vivax* populations are one of the main sources of the genetic make-up of American *P. vivax*. Conversely, the second best scenario (scenario 4: 14.4% of votes) suggested that Ebro could be a migrant from Latin America (Fig. 5a). For this study, the hypothesis by Van Dorp *et al.* on the European origin of *P. vivax* [15] was considered the most plausible.

However, it is important to note that future investigations of the now extinct European gene pool will require additional samples, particularly from the Mediterranean region. Our ABC-RF scenario testing analysis supported the major contribution of European populations in shaping the genetic ancestry of Latin American *P. vivax* populations. Yet, these results also suggest that the colonization history of the Latin American continent might have been more complicated. Specifically, our analysis indicates a potential contribution from West African populations and/or from an unsampled population to the origins of *P. vivax* in Latin America. According to the best supported scenario (scenario 6 in Fig. 5), the West African, Ebro and unsampled populations would have all split almost at the same time. Then, the admixture between the European population and the unsampled population would have composed the Latin American populations. To ascertain the role of West African *P. vivax* as a genetic source in the colonization of Latin America, it is imperative to include additional European *P. vivax* samples from ancient specimens in future studies, and also to increase the West African sample size and it distribution. Furthermore, more elaborated simulation frameworks using demographic-genetic scenarios including gene flow (and not just admixture events) and considering dormancy may bring new insights into the complicated *P. vivax* colonization history of Latin America.

### *P. vivax* probably colonized Latin America more recently than expected

The results obtained in this study showed that European, West African, and unsampled populations all diverged at the same time, about 400 years ago (Fig. 5b). There is no way to determine from where, following divergence, this unsampled lineage originated. This unsampled population could be from Latin America, thus suggesting at least two introductions, 200 years apart, in the 16^th^ century and then in the 19^th^ century in the American continent. Alternatively, this unsampled population could be from West Africa. This scenario is partially supported by the observed shared ancestry with some American populations (Fig. 2c) and our DIYABC-RF analyses. Indeed, in scenarios 4 and 8 (the second and third most supported scenarios in the DIYABC-RF analysis) (Fig. 5a), American populations came from West Africa or from a population descended from West Africa and a European population. Lastly, the unsampled lineage could have been located in North America or the Caribbean where *P. vivax* malaria widely circulated in the latter half of the 19^th^ century [58,59]. Currently, the only way to identify the exact genetic origin of this unsampled population is to generate genetic data from these regions using ancient samples (*e.g.* historical slides, historical mosquito specimens).

The second significant event in the colonization scenario is the admixture between the European population and the unsampled population in which the European population contributed approximately 60% of the current American genetic pool. Our analyses suggest that this event occurred towards the end of the transatlantic slave trade, in the latter half of the 19^th^ century. This timeframe coincides with the massive post-colonial migrations [60], when more than 10 million Europeans settled in Latin America [61], potentially introducing parasites in this continent. It is worth noting that our time estimates are approximately 400 years younger than those estimated by van Dorp *et al.* [15]. Such a difference may be explained by the different methodologies used in the two studies. We used a generation time of 5.5 generations per year [29], but other generation times have been suggested [62] and can also vary depending on the region [63], thus affecting the estimates of divergence times in years. van Dorp *et al.* [15] did not use any generation time estimate and inferred the divergence times by relying on the correlation between collection times and root-to-tip distances of the different samples [64]. This correlation was calibrated using only two historical points: the Ebro composite sample from the 1940s and a North Korean sample from 1953. Furthermore, the fact that Ebro is a composite sample may bias the divergence time estimate. Indeed, the samples in Ebro were collected between 1942 and 1944 and this was not taken into account in the correlation. Moreover, the mutations observed correspond to several isolates, and not to a single one. This may lead to a higher divergence time due to the artificial accumulation of mutations. Unlike van Dorp *et al.* [15], the two analyses used in our study converged toward very similar estimates of the divergence time between Ebro and American populations. Indeed, *Relate* gave a divergence time of 100-300 ybp (Fig. 4c) and DYABC-RF suggested a divergence time of 212 ybp (Fig. 5b).

The approaches used by van Dorp *et al.* [15] and in the present study rely on the Wright-Fisher model for coalescent simulation, which assumes a panmictic population with non-overlapping generations and neutral evolution [65,66]. However, in the case of *P. vivax*, the assumption of non-overlapping generation is not met, particularly because of dormancy. Especially since *P. vivax* emerges from dormancy when the host is infected with new strains [3,67]. As a result, diversity is maintained, and the effective population size is overestimated and the effect of drift is attenuated [57,68]. This may also explain why the Tajima’s D values (S9 Fig.) and population size inference of the American populations (Fig. 4a) did not suggest any bottleneck resulting from colonization, as previously observed [54].

Additionally, our analysis overlooked migrations between populations, that can influence the estimates of genetic parameters, such as divergence times and population sizes. Linked selection may also have biased our inferences [69]. Omitting migration and overlapping generation can reduce the divergence times estilates and increase the inferred population sizes. Linked selection can lead to underestimated the effective population size and divergence times. Incorporating *P. vivax-*specific life-history traits and selection could improve accuracy, but would require long and complicated forward genetic simulations, such as those implemented in *SLiM* 3 [70].

## Conclusion

This study indicates that *P. vivax* invasion of Latin America occurred towards the second half of the 19^th^ century, with a predominantly European contribution. Therefore, *P. vivax* would have been introduced into Latin America by the millions of European settlers during the post-colonial migration. However, Europe was not the only genetic source: at least another population, not currently sampled, might have contributed to the American *P. vivax*. Future studies should focus on obtaining *P. vivax* samples from different parts of the world to identify this population, including also historical samples from Europe, North America, the Caribbean and West Africa.

It is important to note that significant human migrations played a pivotal role in the invasion of the Americas by the two main human malaria agents, *P. falciparum* and *P. vivax*. *P. falciparum* entered the continent with the millions of African slaves who were brought to the Americas from the 16^th^ to the 19^th^ century [17,18], and *P. vivax* might have accompanied millions of European migrants in the late 19^th^ century.

It is now imperative to investigate traces of selection in American *P. vivax* genomes. Understanding the genetic basis of *P. vivax* colonization success in the Americas is crucial for developing effective control strategies and predicting invasions of other regions. Yet, the current literature lacks comprehensive insights into this aspect [71], highlighting the need of more studies in this field.

## Materials and Methods

### *P. vivax* sample collection and ethics statements

In this study, 215 new *P. vivax* isolates from 24 countries were sequenced (Fig. 1). *P. vivax* infection was detected by microscopy analysis, polymerase chain reaction (PCR) amplification of the *Cytochrome b* gene, and/or rapid diagnostic testing. Samples were obtained from patients infected with *P. vivax* following their informed consent and approval from the local institutional review board in each country. The informed consent process for the study involved presenting the study objectives to the community and inviting adults to participate. Before sample collection, each individual was informed about the study purpose and design and provided with a study information sheet. Oral informed consent was obtained from each participant. As the study posed no harm, it did not require written consent for its procedures. This oral consent approach aligned with the ethical standards of each country at the time of enrollment and was endorsed by the local ethics committees. Additionally, measures were taken to ensure the privacy and confidentiality of the collected data through sample anonymization before the study start.

For samples from Honduras, ethical and scientific approvals were obtained from the Ethics Review Committee of the Infectious and Zoonotic Diseases Master’s Program at UNAH (CEI-MEIZ 02–2014; 5/19/2014). Data were analyzed anonymously because the study made secondary use of biological specimens originally collected for malaria diagnosis as per the standard of care in Honduras. For samples collected in Venezuela, each patient gave a written informed consent, and ethical clearance was obtained from the Comité Ético Científico del Instituto de Medicina Tropical de la Universidad Central de Venezuela. Ecuadorian samples were collected through the Ministry of Health malaria surveillance program. The protocol was approved by the Ethical Review Committee of Pontificia Universidad Católica del Ecuador (Approval Number: CBE-016-2013). Informed written consent was obtained from all participants and/or their legal guardians in the case of minors before sample collection. For four samples from Mauritania, the study was approved by the pediatric services of the National Hospital, the Cheikh Zayed Hospital, and the Direction régionale à l’Action Sanitaire de Nouakchott (DRAS)/Ministry of Health in Mauritania. No ethics approval number was obtained at this time. For samples from Azerbaijan and Turkey, *P. vivax* was isolated from patients as part of the routine primary diagnosis and post-treatment follow-up, without unnecessary invasive procedures. The informed consent of each patient or an adult guardian of children enrolled in this study was obtained at the moment of blood collection and included information that samples would be used to investigate the genetic diversity of *Plasmodium* parasites (VIVANIS project, supported by the COPERNICUS-2 RTD project contract ICA2-CT-2000-10046 of the European Commission). For the remaining newly sequenced samples, no specific consent was required because, in coordination with the Santé Publique France organization for malaria care and surveillance, the human clinical, epidemiological, and biological data were collected in the French Reference National Center for Malaria (CNRP) database and analyzed in accordance with the public health mission of all French National Reference Centers. The study of the biological samples obtained in the context of medical care was considered as non-interventional research (article L1221-1.1 of the French public health code) and only required the patient’s non-opposition during sampling (article L1211-2 of the French public health code).

### DNA extraction and sequencing

Genomic DNA was extracted from each sample using the DNeasy Blood and Tissue Kit (Qiagen, France) according to the manufacturer’s recommendations. *P. vivax-like* samples were identified by amplifying with nested PCR the *Plasmodium cytochrome b*, as described in Prugnolle *et al.* [72]. Then, genomic DNA was analyzed on agarose gel (1.8%) in the TAE buffer. Gel-positive amplicons were then sent for sequencing.

For samples identified as *P. vivax*, selective whole-genome amplification (sWGA) was used to enrich submicroscopic DNA levels, as described in Cowell *et al.* [73]. This technique preferentially amplifies *P. vivax* genomes from a set of target DNAs and avoids host DNA contamination. For each sample, DNA amplification was carried out using the strand-displacing phi29 DNA polymerase and *P. vivax*-specific primers that target short (6 to 12 nucleotides) motifs commonly found in the *P. vivax* genome (PvSet1 [73,74]), 30 ng of input DNA was added to a 50-µL reaction mixture containing 3.5 μM of each sWGA primer, 30 U of phi29 DNA polymerase enzyme (New England Biolabs), 1× phi29 buffer (New England Biolabs), 4 mM deoxynucleoside triphosphates (Invitrogen), 1% bovine serum albumin, and sterile water. DNA amplifications were carried out in a thermal cycler with the following program: a ramp down from 35° to 30°C (10 min per degree), 16 hours at 30°C, 10 min at 65°C, and hold at 4°C. For each sample, the products of the two amplifications (one per primer set) were purified with AMPure XP beads (Beckman Coulter) at a 1:1 ratio according to the manufacturer’s recommendations and pooled at equimolar concentrations. Samples with the highest concentration of parasite genome were selected after qPCR analysis using a Light Cycler 96 with the following program: 95°C for 10 minutes; 40 cycles of 95°C for 15 seconds, 60°C for 20 seconds and 72°C for 20 seconds; 95°C for 10 seconds; and 55°C for 1 minute. Last, each sWGA library was prepared using the two pooled amplification products and a Nextera XT DNA kit (Illumina) according to the manufacturer’s protocol. Each sWGA library was prepared using the two pooled amplification products and a Nextera XT DNA kit (Illumina) according to the manufacturer’s protocol. Samples were then pooled and clustered on a Hiseq 350M package sequencer or Novaseq S4 1 lane PE150 with 2 × 150-bp paired-end reads.

### Combining newly sequenced and literature genomic data

The newly sequenced dataset for this study included 74 American *P. vivax* isolates (3 from Brazil, 19 from Ecuador, 22 from French Guiana, 4 from Guyana, 9 from Honduras, and 17 from Venezuela), 63 African isolates (1 from Central Africa, 21 from Comoros, 9 from Djibouti, 1 from Ethiopia, 1 from Gabon, 1 from Mali, 16 from Mauritania, 1 from Mayotte, 8 from Sudan, 2 from Chad, 1 from Uganda, and 1 from an unclear location between Chad and Mali), 40 Middle Eastern isolates (23 from Afghanistan, 4 from Armenia, 11 from Azerbaijan, and 2 from Turkey), 37 South Asian isolates (19 from India, 18 from Pakistan), and 1 Oceanian isolate (PNG).

In addition, genomic data from various sources were added: from Daron *et al.* [29], Benavente *et al.* [28], and the MalariaGEN project *P. vivax* Genome Variation [30]. From the large *P. vivax* Genome Variation dataset of the MalariaGEN project (n=1,895), only samples accessible for which fastq file data were available, passing the quality check defined by Adam *et al.* [30], and that were not related were selected (S1 Fig.). The relatedness between haploid genotype pairs within each country was measured by estimating the pairwise fraction of the IBD between strains within populations using the *hmmIBD* program [75], with default parameters for recombination and genotyping error rates, and using the allele frequencies estimated by the program. Isolate pairs that shared >50% of IBD were considered highly related. In each family of related samples, only the strain with the lowest number of missing data was retained. This left 500 samples from the MalariaGEN *P. vivax* Genome Variation project [30]. As the main focus of this study was on Latin America, samples from Malaysia, Myanmar, Thailand, and Cambodia were not retained because we already had more than 20 samples in each country. In total, 295 samples from the MalariaGen *P. vivax* Genome Variation project [30], 499 samples from Daron *et al.* [29], and 125 samples from Benavente *et al.* [28] were selected. The 499 samples from Daron *et al.* [29] included 26 African great apes *P. vivax-like* samples from three African countries (Cameroon, Gabon, and Ivory Coast).

As van Dorp *et al.* [15] inferred a European origin of the American populations, the ancient Spanish sample (Ebro) from this publication was added. Ebro is a composite sample of four samples, from four different slides dating from the 1940s: CA, POS, CM, and lane-8. Samples were downloaded from ENA (study accession PRJEB30878) and the fastq were single ends. The whole dataset from the literature comprised 920 samples that were added to the 214 newly sequenced genomes.

### *P. vivax* and *P. vivax-like* read mapping and SNP calling

The *P. vivax* and *P. vivax-like* read mapping and SNP calling steps are detailed in S3 Figure. All modern sample sequencing reads were trimmed to remove adapters and preprocessed to eliminate low-quality reads (--quality-cutoff=30) using *cutadapt* v1.18 [76]. Reads shorter than 50 bp containing “N” were discarded (--minimum-length=50 --max-n=0). Sequenced reads were aligned to the *P. vivax* reference genome PVP01 [77] with *bwa-mem* v0.7.17 [78].

For Ebro, a different pipeline was used to consider the specific features of ancient DNA. The 3’ adapters, reads with low-quality score (--quality-cutoff=30), and reads shorter than 30 bp were removed with *cuadapt* [76]. Then the cleaned fastq were merged, to create the composite sample as described in van Dorp *et al.* [15]. As the European samples were coinfected by *P. falciparum*, all reads were mapped to the *P. vivax* reference genome PVP01 [77] and to the *P. falciparum* reference genome Pf3D7 v3 [79] using *bwa-mem* [78]. For reads that mapped to both reference genomes, the edit distance was extracted and reads that aligned equally or better with *P. falciparum* than with *P. vivax* were removed. Then reads with mapping quality <30 and duplicates were removed with *samtools* v1.9 [80] and with *Picard tools* v2.5.0 (<u>broadinstitute.github.io/picard/</u>), respectively. Then, the base quality was rescaled with *MapDamage 2* [81,82] to account for postmortem damage at the read ends.

For all samples, the *Genome Analysis Toolkit* (*GATK* v3.8.0 [83]) was used to call SNPs in each isolate following the *GATK* best practices. Duplicate reads were marked using MarkDuplicates from the *Picard tools* with default options. Local realignment around indels was performed using the IndelRealigner tool from *GATK*. Variants were called using the HaplotypeCaller module in *GATK* and reads mapped with a “reads minimum mapping quality” of 30 and minimum base quality of >20. During SNP calling, the genotype information was kept for all sites (variants and invariant sites, option ERC) to retain the information carried by the SNPs fixed for the reference allele. Then only the nuclear genome was kept, filtered out with *BCFtools* v1.10.2 [80,84], and all individual VCF files were merged with *GATK*. Organelle genomes (mitochondria and apicoplast) were not included because they are haploid markers, without recombination and with only maternal heritability [85–87].

### Data filtering

Combining data from the literature (n=919) with the 214 newly sequenced samples resulted in a total of 1,133 samples for the unfiltered dataset: 1,106 modern human *P. vivax* samples, 26 modern African great apes *P. vivax-like* samples, and one ancient human *P. vivax* sample. All filtration steps are detailed in S1 Figure.

For all modern samples, samples with >50% missing data were removed. As several strains can infect the same host, the within-host infection complexity was assessed with the *F*_WS_ metric [88], calculated with *vcfdo* (github.com/IDEELResearch/vcfdo; last accessed July 2022). Samples with mono-clonal infections, *i.e. F*_WS_ >0.95 were kept. Highly related samples and clones could have generated spurious signals of population structure, biased estimators of population genetic variation, and violated the assumptions of the model-based population genetic approaches used in this study [89]. The relatedness between haploid genotype pairs was measured by estimating the pairwise fraction of the genome IBD as explained above. Within each country, isolate pairs that shared >50% of IBD were considered highly related. In each family of related samples, only the strain with the lowest number of missing data was retained. The final dataset included 619 modern *P. vivax* individuals from 34 countries, two *P. vivax-like* from Cameroon, infecting Nigeria-Cameroon chimpanzees (*Pan troglodytes ellioti*), and Ebro, an ancient DNA sample from Spain.

Two dataset formats were used in function of the analysis: (*i*) a VCF containing all the samples, and (*ii*) bam files of the samples for the *ANGSD* v0.940 [31] or *ANGSD* suite software (S1 Fig.). For both formats, the analysis focused only on the core genome regions as defined by Daron *et al.* [29].

For the VCF, only bi-allelic SNPs with ≤20% missing data and a quality score ≥30 were kept. The minimum and maximum coverage were 10 and 106. Moreover, singletons were removed to minimize sequencing errors. Finally, 950,034 SNPs with a mean density of 41.14 SNPs per kilobase were retained.

For the *ANGSD* suite software, all sites with a base quality ≥20, a maximum total depth of 106 X, a minimum total depth of 5 X, and present in at least five individuals were kept. For analyses that did not require invariant sites, only bi-allelic SNPs with a *p-value* = 1e-6 were retained. In total 20,844,131 sites from the core genome were considered, including 1,437,273 variants (69.59 SNPs/kb).

### Population structure

The PCA and the ancestry plots were computed with *ANGSD* and *PCAngsd* v0.98, after selecting only biallelic SNPs present in the core region of the *P. vivax* genomes and excluding SNPs with a minor allele frequency (MAF) ≤5%. The variants were LD-pruned to obtain a set of unlinked variants using *ngsld* v1.1.1 [90], with a threshold of r²=0.5 with windows of 0.5kb. In total, 105,527 SNPs were used for 620 individuals (*P. vivax-like* samples were removed). For the ancestry plots, obtained from *PCAngsd*, the k value (number of clusters) was from 2 to 16. Then, *pong* v1.5 [91] was used to analyze the *PCAngsd* outputs and compute ancestry proportions.

The ML tree was obtained with *IQ-TREE* v2.0.3 [33] using the best-fitted model determined by *ModelFinder* [35]. As the dataset comprised 950,034 SNPs of the core genome and no constant site, the ascertainment bias correction was added to the tested models. The best inferred model was a general time reversible (GTR) model of nucleotide evolution that integrated unequal rates and unequal base frequency. The node reliability was assessed with Ultrafast Bootstrap Approximation [92] and the SH-aLRT test [93].

### Relationships between populations

Population networks were estimated using two different approaches: *TreeMix* v1.13 [40] and *AdmixtureBayes* (github.com/avaughn271/AdmixtureBayes; last accessed June 2023) [41]. Unlike with the “classic” ML phylogenetic tree at the level of individual strain genomes (see above), these two methods use allele frequency (co-)variation within and among populations to derive the fraction of shared versus private genetic ancestry (or genetic drift) and to infer the population branching order, while taking into account population genetic processes, such as genetic drift, historical migration, and admixture events between populations [94,95].

To be able to compare results, the same dataset was used for both approaches: the VCF dataset was restricted to SNPs without missing data in Ebro (17,696 variants) and to subsampled modern populations as follows. Only populations with the largest sample sizes for each genetic cluster (according to PCA and ancestry plots) were kept, as well as all the American populations. The SNP data set was LD-pruned with With *PLINK* v2 [96], a threshold of r²=0.5, with sliding windows of 50 SNPs and a step of 10 SNPs, and MAF filtering with a threshold of 5% (9,022 SNPs remaining). In some populations, there were still positions with missing data, and those positions were removed because *TreeMix* do not handle missing data. This filtering resulted in 7,109 SNPs for 18 ingroup populations (from n=1 to n=80) and one outgroup population (*P. vivax-like*).

For the *TreeMix* analyses, the number of migration events (*m*) that best fitted the data were estimated by running *TreeMix* 15 times for each *m* value, with *m* ranging from 1 to 10. The optimal *m* value was estimated using the *OptM* R package [97]. Then, a consensus tree with bootstrap node support was obtained by running *TreeMix* 100 times using the optimal *m* value. Results were post-processed using the *BITE* R package [98].

For the *AdmixtureBayes* analyses, three independent runs were performed, each including 40 Markov Chain Models and 250,000 steps, to identify convergence. Convergence was checked with a Gelman-Rubin Plot, as recommended by Nielsen *et al*. [41]. This analysis found that two chains had converged (S7 Fig.). The results of these two chains were analyzed after a burn-in of 50% and a thinning step of 10, using the default parameters. The common network among the three best networks in each chain that converged was retained, based on posterior probabilities.

From the dataset generated with *PLINK* for *TreeMix* and *AdmixtureBayes*, the *f_3_*-statistics were calculated using *ADMIXTOOLS 2* v2.0.0 [99].

### Demographic and invasion history of *P. vivax* in the Americas

Tajima’s D [44] and nucleotide diversity (π) were measured for each population with ≥10 individuals to avoid biases [100]. The sample size was standardized at 10 (*i.e.*, 10 randomly chosen isolates for each population) to obtain values that could be compared. Tajima’s D and nucleotide diversity (π) values were estimated using *ANGSD* [31], with a window of 500 bp and a step of 500 bp in the core genome, and only windows with ≥50 sites were kept.

The coalescent-based approach of *Relate* [45] and the core region of the genome of all chromosomes were used to infer historical changes in *N_e_* and to estimate the divergence time between modern populations. *Collate* was used to calculate CCR between Ebro and modern populations, because this program was designed to handle ancient DNA samples with low coverage [46]. For both analyses, a mutation rate of 6.43 x 10^-9^ mutations/site/generation [29] and a generation time of 5.5 generations/years [29] were used. Alleles were polarized using the two *P. vivax-like* samples as an outgroup.

A genetic map was required to carry out these analyses. This was generated using *LDhat* [101] and considering the Thailand population, on one of our most diverse populations. First, individuals with ≥5% missing data were removed, leaving 58 samples. The function *pairwise* was used to estimate a first approximation of Watterson’s Theta and then, the likelihood table was estimated with these parameters: *-n 58 -rhomax 100 -n_pts 101 -theta 0.00011 -split 8*. The recombination map was then obtained by using the function *interval* (-*its 1100000 -samp 100 -bpen 5*) followed by *stat* with a burn-in of 10,000 samples. The recombination map (4N_e_r/pb) was co,verted into a genetic map (cM/Mb) with a *N_e_* estimate of 6,000 (inferred with *Stairwayplot 2* on chromosome 14).

### Scenarios for *P. vivax* invasion of the Americas

Different possible scenarios of population divergence and admixture were evaluated using *DIYABC Random Forest* (RF) [48] via the *diyabcGUI* v1.2.1 R package.

As Ebro is a composite sample, it was made haploid by creating two haploid samples, splitting the “heterozygous” sites between the two samples. To minimize the effect of missing data, the sites genotyped in at least one individual in each population and at least in 40% of individuals in all populations combined were kept. Moreover, individuals with ≥ 50% missing data were removed. Thus, 45 haploid individuals for three populations and 3,139 SNPs were retained.

Twelve scenarios were tested: (1) American populations as a result of West African and European admixture; (2) uniquely European origin; (3) uniquely West African origin; (4) Ebro as a migrant from Latin America with American populations descending from West African ancestors; (5) Ebro resulting from the admixture between American populations and an unsampled West African “ghost” population; the American populations would have split from an ancestral population; (6) American populations stemming from the admixture of European and unsampled populations; (7) American populations from Europe with a secondary West African contribution; (8) American populations from West Africa with a secondary European contribution; (9) American populations from Africa with two introduction waves; (10) American populations originating from Europe, with Ebro resulting from an American-European ghost population admixture; (11) American populations resulting from two successive admixture events, the first and oldest one between the European populations and an unsampled ghost population, followed by a more recent wave of introduction from West Africa; and (12) Ebro resulting from the American-ghost population admixture, with American populations splitting from West Africa.

The scenario parameters were considered as random variables drawn from prior uniform distributions (S3 Table). *DIYABC-RF* was used to simulate 20,000 genetic datasets per scenario with the same properties as the observed data set (number of loci and proportion of missing data). Simulated and observed datasets were summarized using the whole set of summary statistics proposed by *DIYABC-RF* for SNP markers to describe the genetic variation of each population (e.g., proportion of monomorphic loci, heterozygosity), pair of populations (e.g., *F*_ST_ and Nei’s distances), or trio of populations (e.g., *f*_3_-statistics, coefficient of admixture). The total number of summary statistics was 50.

Then with these 20,000 simulated data sets per scenario, the *RF* classification procedure was used to compare the likelihood of the competing scenarios. *RF* is a machine-learning algorithm that uses hundreds of bootstrapped decision trees to perform classifications, using the summary statistics as a set of predictor variables. Then a classification forest of 1500 trees was grown.

Some simulations were excluded from the decision tree building during each bootstrap (*i.e.*, out-of-bag simulations). Instead, they were used to calculate the prior error rate and the type II error rate. Additionally, the confusion matrix was computed to provide a more global assessment of the RF procedure performance [102] (S10 Fig.). The result convergence was evaluated over ten independent RF runs, as recommended by Fraimout *et al.* [103]. To determine the compatibility of the formulated scenarios and associated priors with the observed dataset, we plotted the summary statistics for both on the first two principal axes of a linear discriminant analysis and of a PCA (S10 Fig.), following the recommendation by Pudlo *et al*. [104]. The location of the summary statistics from the observed data within the clouds formed by those from the simulated data was visually checked for at least one scenario (S10 Fig.).

The RF computation provides a classification vote for each scenario (*i.e.*, the number of times a model is selected from the decision trees). The scenario with the highest classification vote was selected as the most likely scenario.

Then, the posterior distribution values for all parameters for the best model identified were estimated using the derivation of a new RF for each component of interest of the parameter vector [105], with classification forests of 1500 decision trees, and based on a training set of 100,000 simulations. Estimated parameters included the effective size (*N_e_*) for each population, split times among populations, and admixture event timing and rates. Estimates for timing parameters were converted from generations to years assuming a generation time of 5.5 generations per year [29].

## Data availability

The raw genomic data used in this study are available on NCBI and NCBI SSR-ID, bioproject, biosample, and sources are indicated in Supplementary Table 1.

The scripts are available in this github repository, as well as the genetic map used for *Relate* and the header file of the *DIYABC-RF* analysis with the scenarios: https://github.com/MargauxLefebvre/LatinAmerica_vivax.

## Acknowledgments

The authors acknowledge the ISO 9001-certified IRD i-Trop HPC (member of the South Green Platform) at IRD Montpellier for providing HPC resources that contributed to the research results reported within this paper. URL: https://bioinfo.ird.fr/- http://www.southgreen.fr. The bioinformatics analyses were also performed on the Core Cluster of the Institut Français de Bioinformatique (IFB) (ANR-11-INBS-0013).

This work was supported by the French National Research Agency (project JCJC GENAD ANR-20-CE35-0003 and MICETRAL ANR-19-CE350010), CNRS, and the French Institute for Public Health Surveillance (Santé publique France) (grant No CNR Paludisme). The collection of samples in Ecuador was funded by Pontificia Universidad Católica del Ecuador (Grants M131416 and N131416).

The authors also thank the members of the French National Reference Centre for Imported Malaria Study Group for providing *P. vivax* isolates. We would also like to thank Clarice Moulin for his help in the laboratory work on the newly sequenced samples, as well as Elisabetta Andermarcher for proofreading the manuscript and supplementary material.

## Author Contributions

Conception: V.R., F.P., M.C.F. and M.J.M.L.; funding acquisition: V.R. and F.P.; biological data acquisition and management: V.R., G.F., O.N, S.H, C.S, B.P, A.B, J-F.T and F.E.S; sequence data acquisition: F.D. and C.A.; method development and data analysis: V.R., M.J.M.L., M.C.F., and F.P.; interpretation of the results: V.R., F.P., M.C.F. and M.J.M.L.; drafting of the manuscript: M.J.M.L., V.R., M.C.F. and F.P.; and reviewing and editing of the manuscript: V.R., F.P., M.C.F. and M.J.M.L.

