## Supplementary Figures and Tables for "Genomic exploration of the complex journey of *Plasmodium vivax* in Latin America"

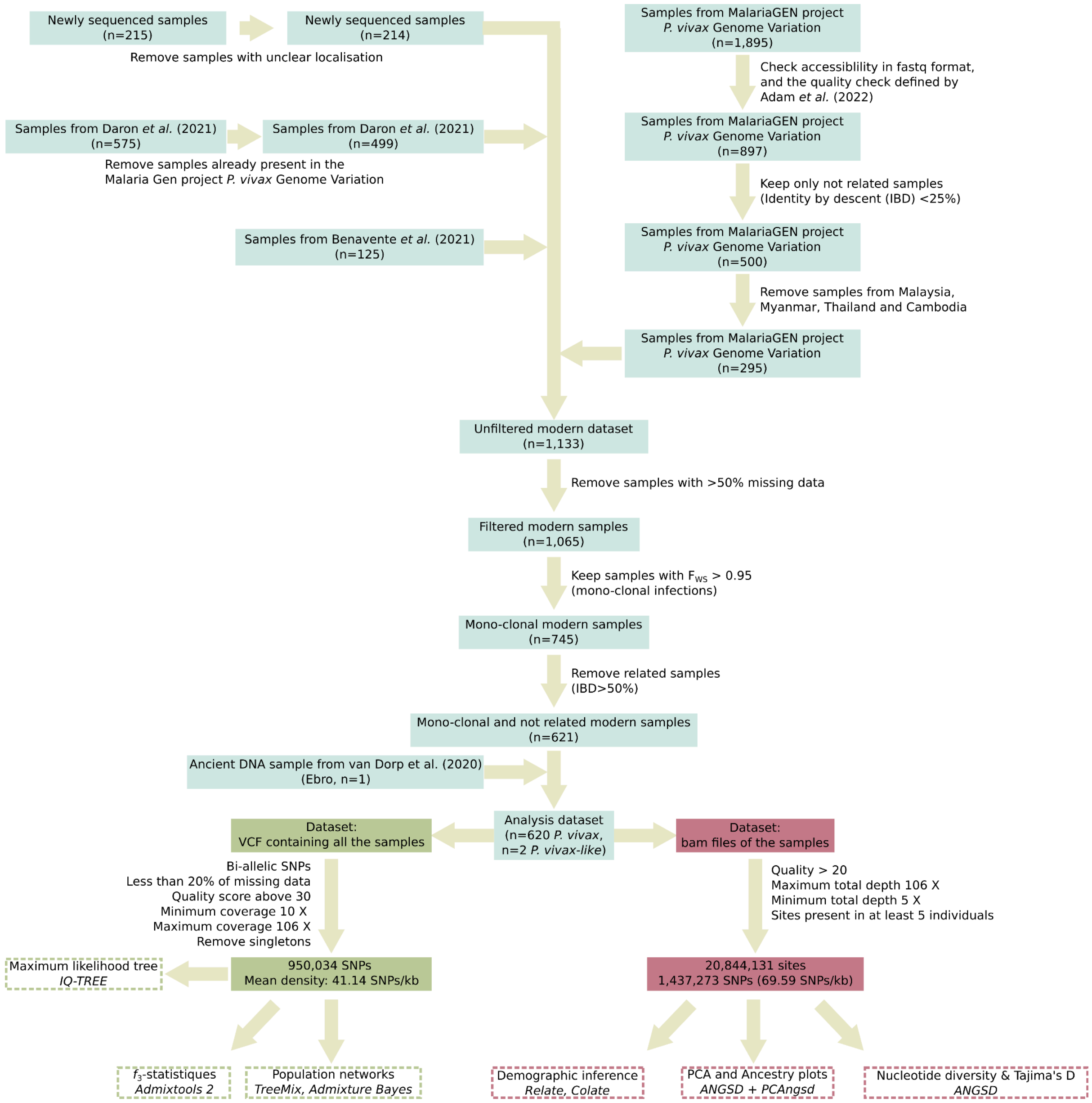


**S1 Figure: Filtering and dataset creation.** Each box in the diagram represents a specific filtering step and also indicates the number of remaining samples or SNPs. Key filtering options also are indicated. The final steps (dotted line boxes) show the analyses performed using the different datasets.

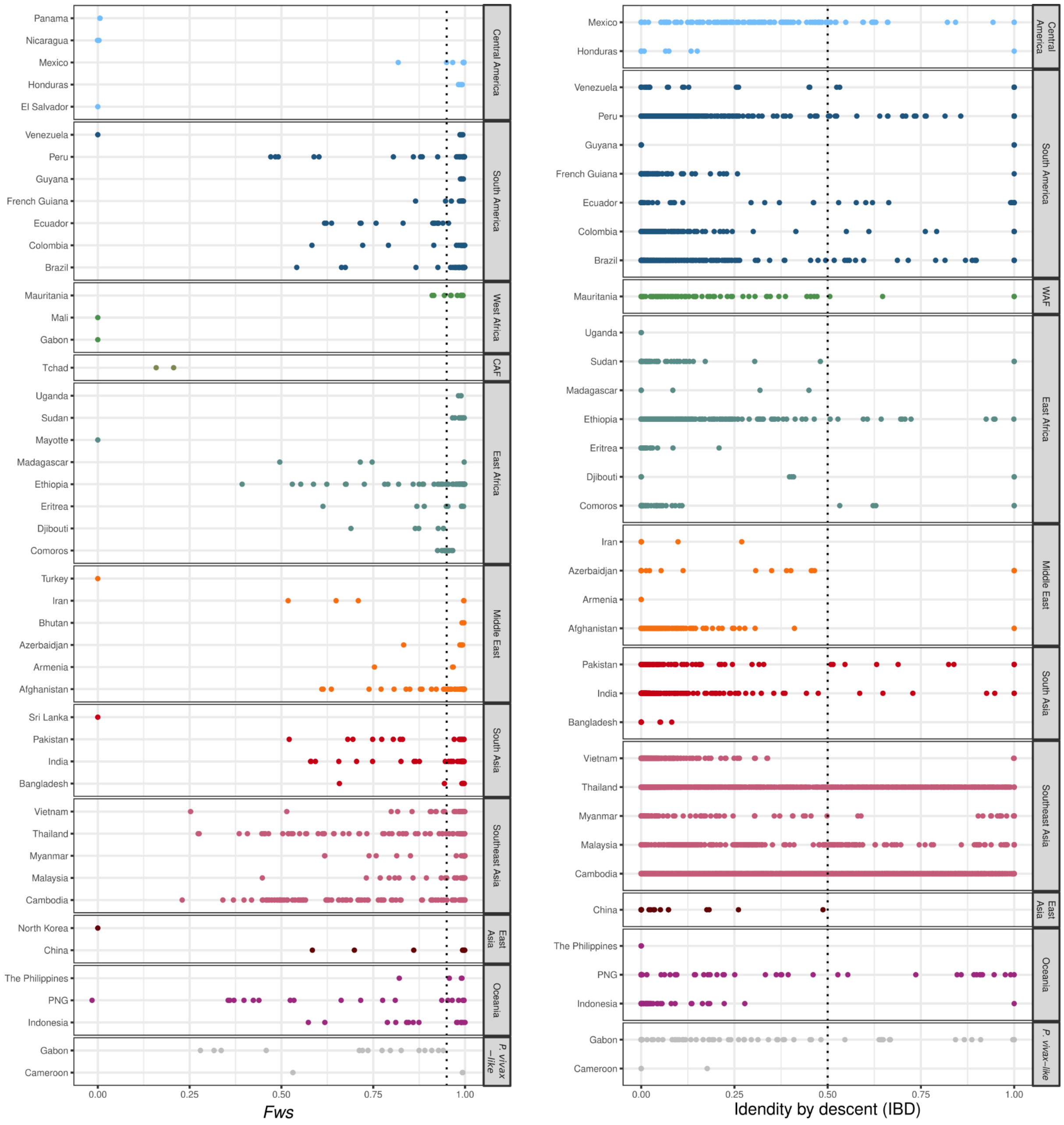


**S2 Figure: Within-sample infection complexity ($F_{WS}$ index) and inbreeding (identity by descent) in *P. vivax* and *P. vivax-like* populations.** The $F_{WS}$ index is a proxy of the diversity within individual infections, from 0 (high diversity) to 1 (no diversity). $F_{WS}$ values >0.95 (indicated by a dotted line) usually indicate monoclonal infections. Identity by descent (IBD) indicates the percentage of the genome resulting from inbreeding among pairs of individuals of the same population. A pairwise IBD >0.5 (dotted line) resulted in the exclusion of one of the two individuals in the strain pair. CAF= Central Africa, WAF= West Africa. PNG=Papua New Guinea.

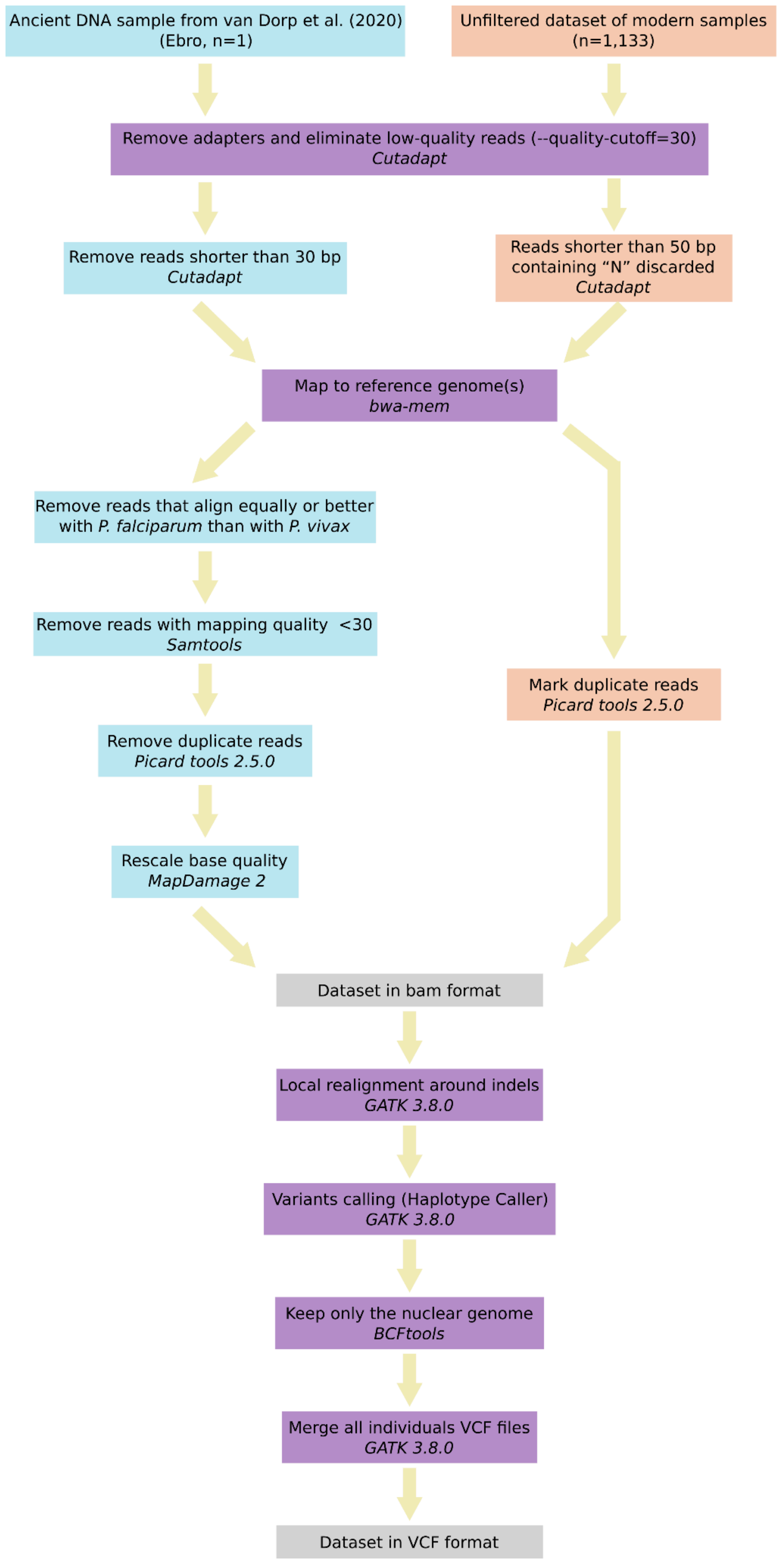


**S3 Figure: *P. vivax* and *P. vivax-like* read mapping and SNP calling steps.** The steps in blue are specific to the ancient DNA Ebro sample, while the steps in red are specific to modern samples. Steps in purple are common to both modern and ancient samples. The gray steps highlight the datasets in the two formats used in this study: bam and VCF.

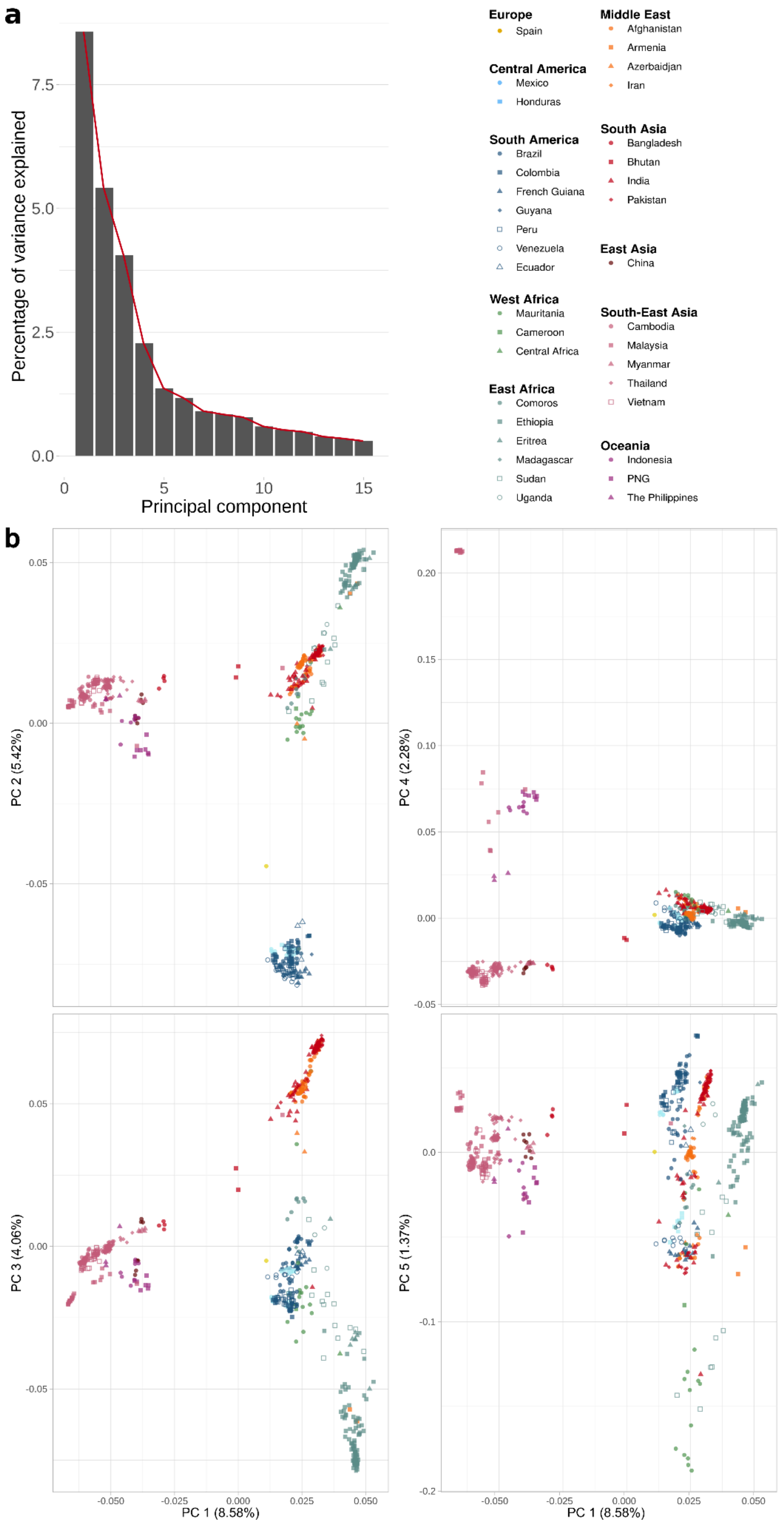


**S4 Figure: Principal component analysis (PCA) for 619 modern *P. vivax* strains and the Ebro ancient DNA sample from Spain based on the genotype likelihood of 105,527 unlinked SNPs. (a)** Percentage of variance explained by the first 15 principal components (PC). The optimal number of PCs is 5, as determined by the elbow (broken-stick) method. **(b)** PCA plots for PC 1 to 5. PNG: Papua New Guinea.

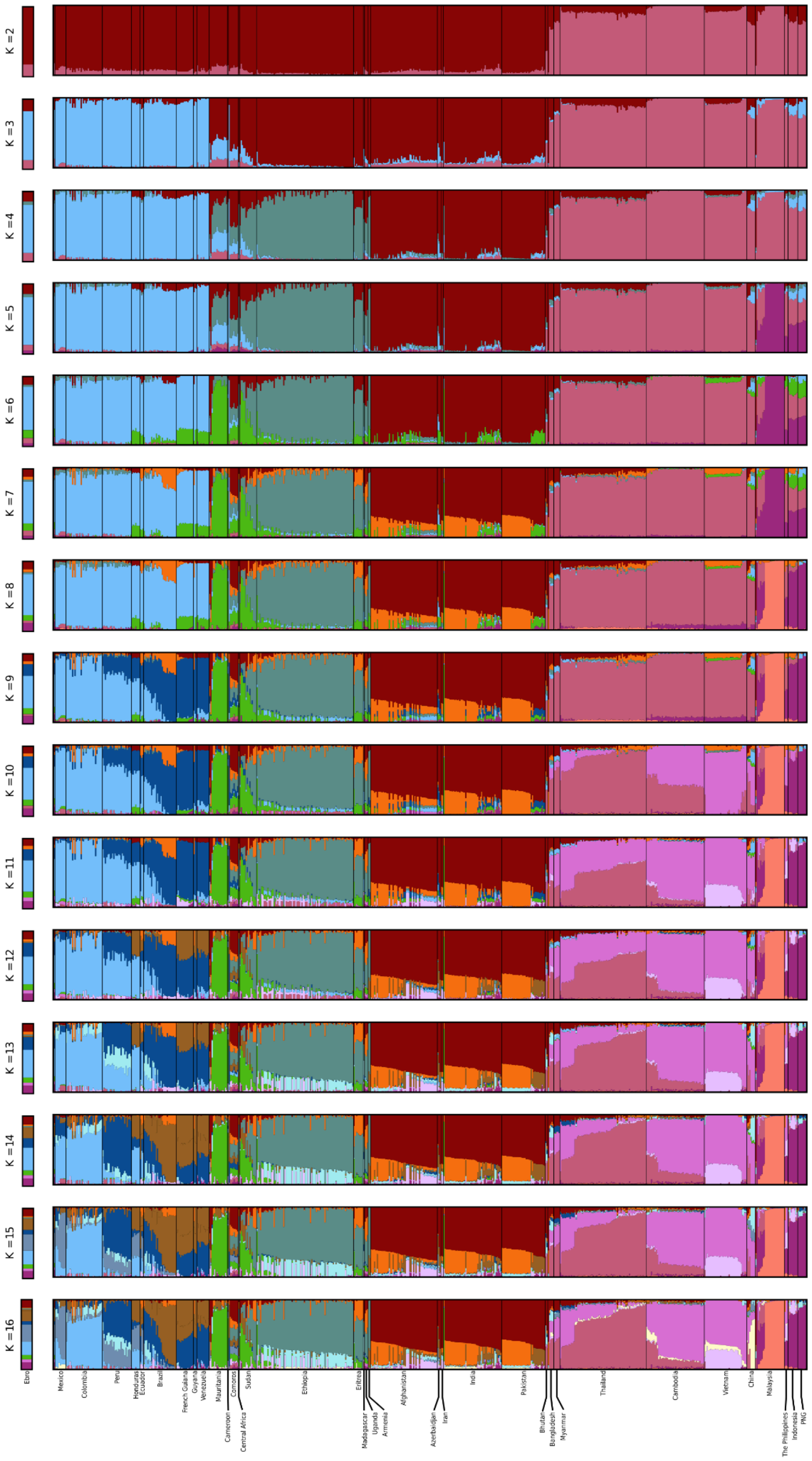


**S5 Figure: Genetic ancestry of *P. vivax* populations worldwide estimated with *PCAngsd* (K=2 to K=16).** The number (K) of clusters tested is specified on the left. According to Meisner and Albrechtsen [1], the best K is determined by 1+ D (the optimal number of principal components). As presented in S4 Figure, D would be = 5 (determined by the elbow (broken-stick) method), thus K=6. PNG: Papua New Guinea.

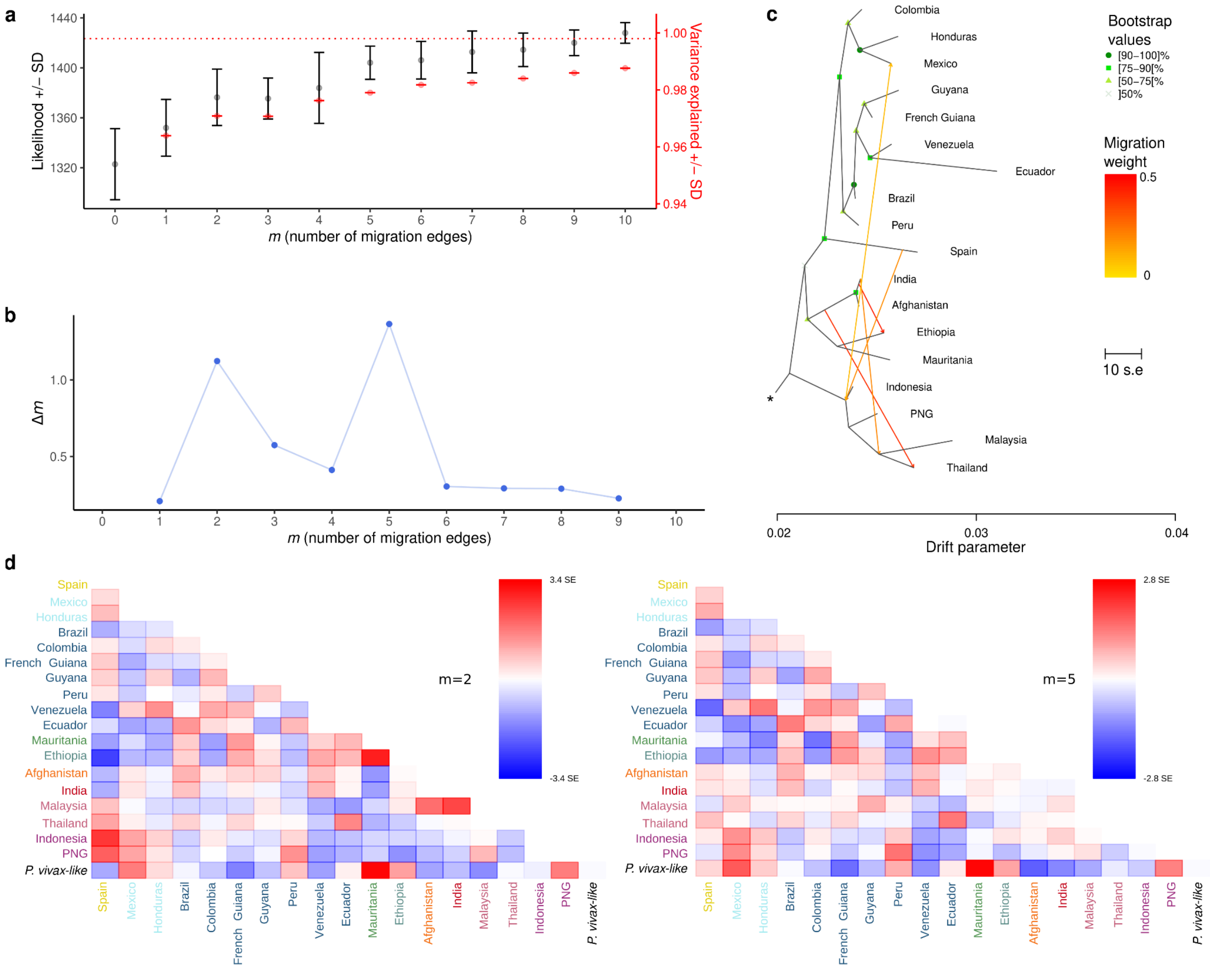


**S6 Figure: Identification of the optimal number of migration edges (*m*) for 18 *P. vivax* populations and consensus topology for m=5 with *TreeMix*.** (**a**) Changes in the mean likelihood score (± SD) and the mean total fraction of the genetic variance explained (± SD) in function of the number of migration edges in the *TreeMix* analysis. (**b**) The second-order rate of change in likelihood (Δ*m*) across migration edges (*m*) values. The *OptM* R package [2] and the Evanno method [3] were used for panels A and B, using 15 replicates for each migration edge (*m)* value, from 0 to 10. The inflection points were observed at *m*=2 and *m*=5. (**c**) *TreeMix* tree of a subset of 18 *P. vivax* populations with five migration edges (arrows), rooted with *P. vivax-like* indicated with the asterisk. The scale bar shows ten times the mean standard error (s.e.). The migration weight is indicated with a color scale, from yellow (0%) to red (50%). (**d**) Visualization of the residuals from the fit of the model to the data for the trees with *m*=2 and *m*=5. The color scales indicate the residuals in standard error units, red for positive values and blue for negative values. PNG: Papua New Guinea.

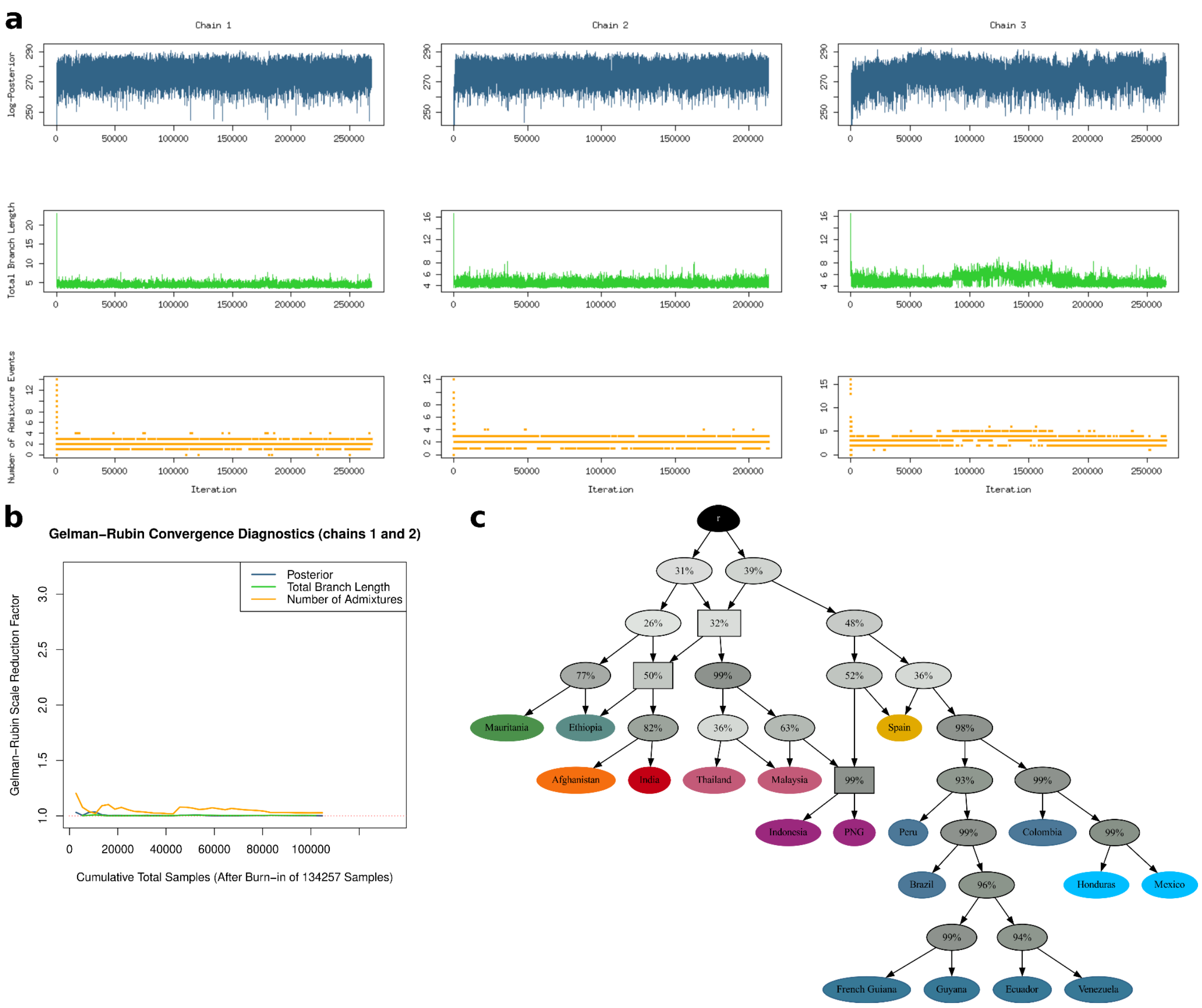


**S7 Figure: Identification of the optimal number of admixture events for 18 *P. vivax* populations and consensus topology with *AdmixtureBayes*. (a)** Trace plots of the posterior probability, total branch length, and number of admixture events for each chain. **(b)** The plot of the Gelman-Rubin convergence diagnostics on chains 1 and 2 for three summary statistics after a burn-in fraction of 50%. The rapid convergence to 1 indicates that this is a sufficient burn-in period. **(c)** Consensus tree generated by combining nodes with a posterior probability >25% of appearing in the admixture graph. Square nodes represent admixture events. The percentages in the nodes are the posterior probability that the true graph has a node with the same descendants. PNG: Papua New Guinea.

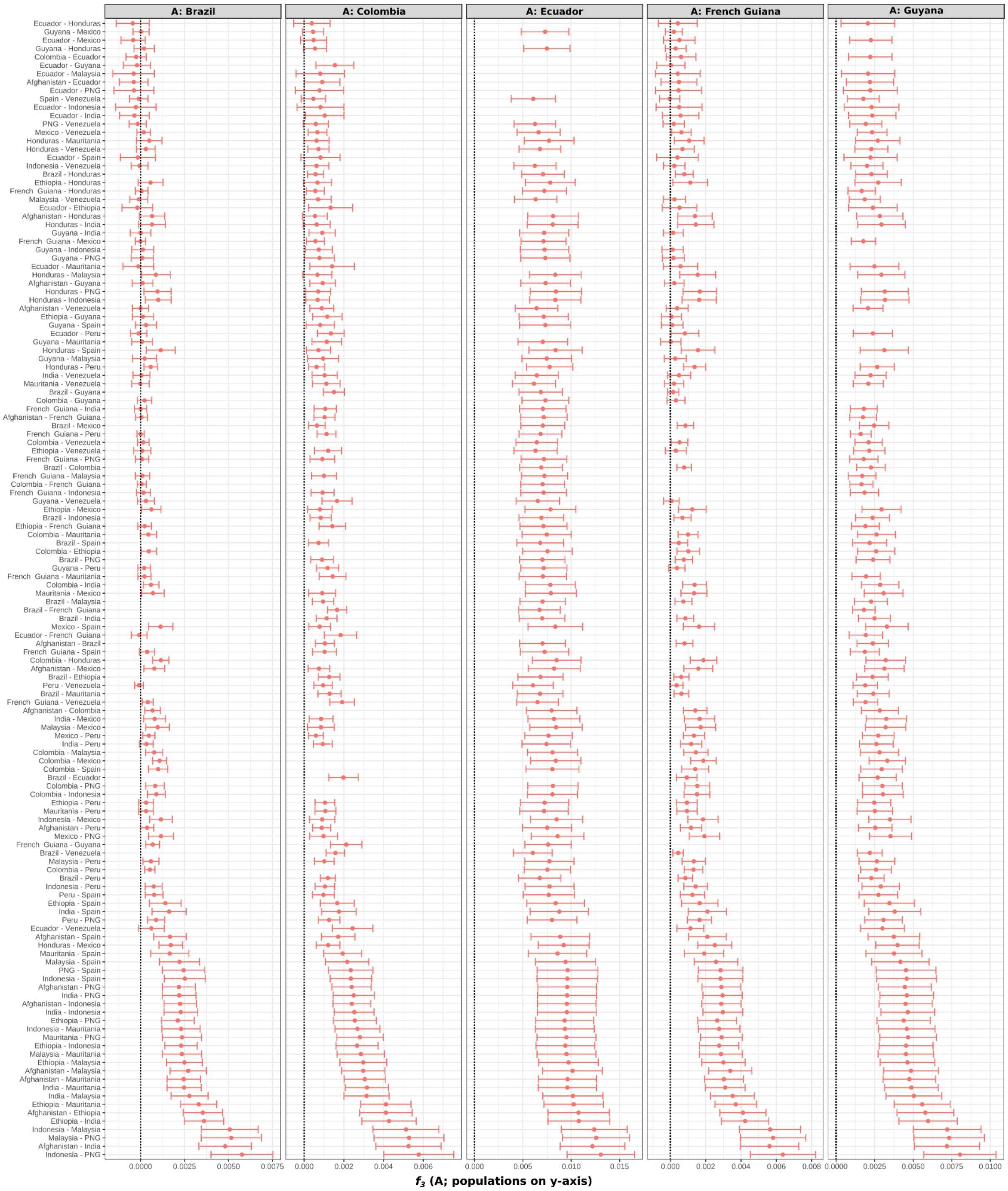


**S8 Figure: $f_3$-statistics for American populations with possible source populations from other American populations and other *P. vivax* populations of the world (Africa, Asia, and Europe).** The dotted line marks the zero. All values are not significantly negative. PNG: Papua New Guinea.

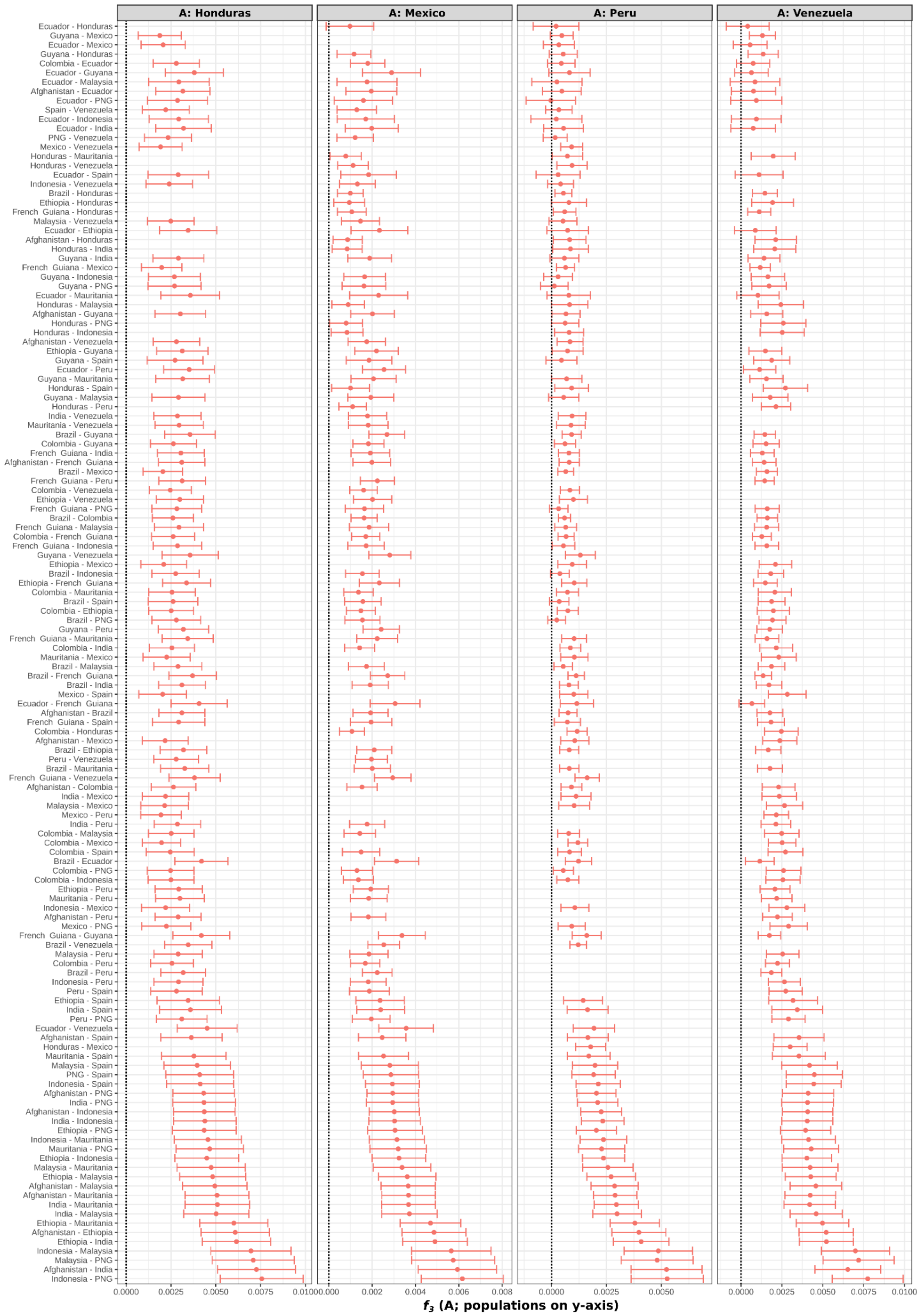


**S8 Figure (continued): $f_3$-statistics for American populations with possible source populations from other American populations and other *P. vivax* populations of the world (Africa, Asia, and Europe).** The dotted line marks the zero. All values are not significantly negative. PNG: Papua New Guinea.

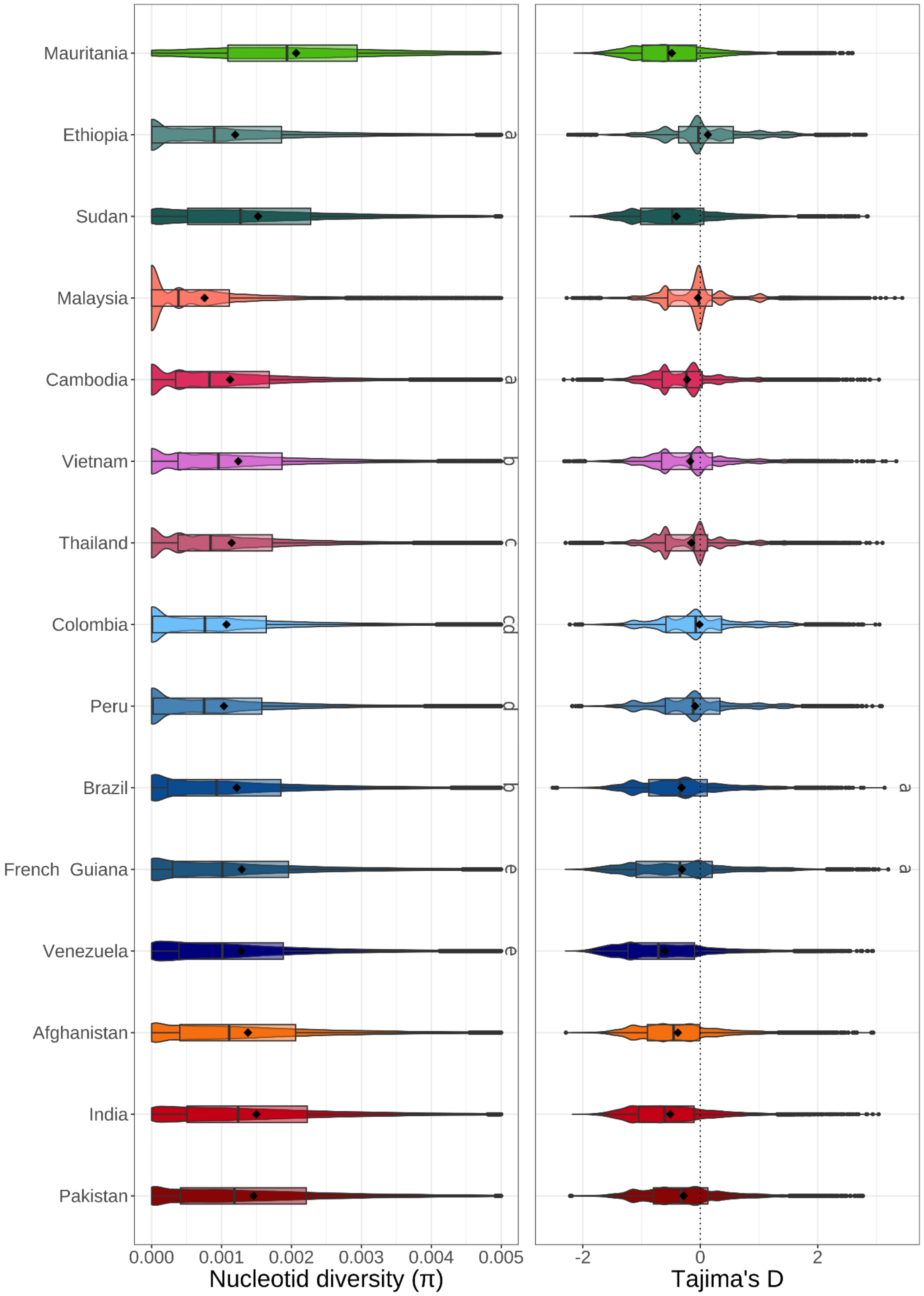


**S9 Figure: Genetic diversity of *P. vivax* populations.** Distributions of the nucleotide diversity (π) and Tajima's D values for populations with >20 samples. The same superscript letters indicate no significant difference between the distributions (Wilcoxon test and Bonferroni correction). If there is no superscript, the distribution is significantly different from all other distributions with a *p-value* < 0.003.

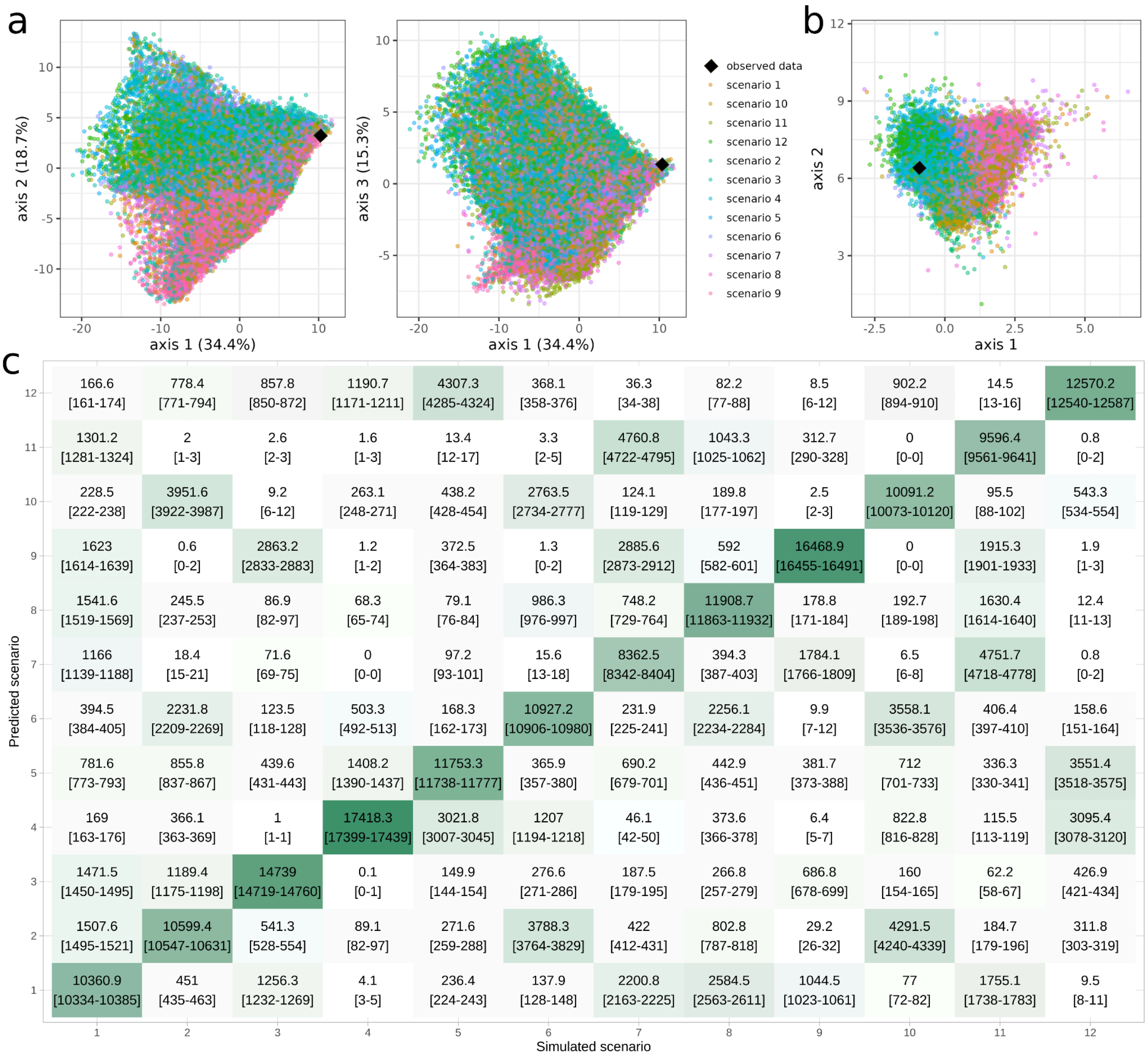


**S10 Figure: ABC-RF performance to select the best model. (a)** Principal Component Analysis (PCA) based on the summary statistics generated by the simulated datasets from the training set (a color indicates each simulated model) and the observed dataset (indicated with a black diamond). **(b)** Projection of the reference table on the first two Linear Discriminant Analysis (LDA) axes. Colors correspond to the model indices. A black diamond indicates the location of the observed dataset. **(c)** ABC-RF out-of-bag confusion matrix. The diagonal represents the proportion of simulated datasets correctly classified for each demographic scenario. Each column corresponds to the scenario under which simulations were generated and each row is the best-supported scenario selected by the ABC-RF classifier. The values represent the mean number of votes over 10 replicates, also represented by the color gradient (the darker the color, the more votes). The minimum and maximum number of votes is indicated between brackets.

**S1 Table: Samples metadata and listing.**
The NCBI SSR-ID, bioproject, biosample, and source are indicated for each sample. When available, the latitude and longitude are specified. NA, information not available. The QC column indicates whether samples have successfully passed the quality control (QC) and are included in the final dataset for analyses. For samples that did not pass the QC, the reasons are outlined in the "Reasons_QC" column as follows: "Missing data" for samples with >50% missing data, "Low $F_{ws}$" for samples removed due to multi-clonal infections, and "High IBD" for samples that are related to those kept in the analysis dataset. For additional details, please refer to the Materials and Methods section and S2 Figure.

**S2 Table: Metadata for the analysis dataset.**
For each sample, the number of reads and the mean coverage are indicated. Each sample metadata included the percentage of the genome covered by at least 1X (% > 1X), 5X (% > 5X), and 10X (% > 10X) sequencing depth. When available, the latitude and longitude are specified. NA: information not available.

**S3 Table: Distribution and conditions of the DIYABC-RF scenario parameters.** To see the correspondence between the parameters and the scenarios, refer to Fig. 5a. All distributions are uniform.

| Parameter name | Minimum | Maximum | Parameter name | Minimum | Maximum |
|---|---|---|---|---|---|
| ***Population sizes*** | | | ***Event times (in generations)*** | | |
| Nbot | 1 | 10 | tdivadm | 500 | 5,500 |
| Nanc | 100 | 150,000 | tbot | 1 | 10 |
| NCol | 10 | 100,000 | tdivebraf | 500 | 5,500 |
| NMauri | 10 | 10,000 | tdivebram | 500 | 5,500 |
| NEbro | 10 | 50,000 | tdivamaf | 750 | 8,500 |
| NAncEbro | 10 | 50,000 | tbotebro | 1 | 10 |
| NbotEbro | 1 | 10 | tadmebro | 500 | 6,500 |
| Nanc1 | 200 | 150,000 | tdiv1/tdiv1b | 500 | 8,500 |
| Nanc2 | 200 | 150,000 | tdiv2/tdiv2b | 750 | 8,500 |
| Nghost | 200 | 100,000 | tct2 | 750 | 7,500 |
| Nghostam | 10 | 100,000 | tdivamaf2 | 750 | 6,500 |
| NghostEbro | 10 | 50,000 | tct3 | 750 | 7,500 |
| Nghost2 | 10 | 100,000 | ***Conditions*** | | |
| ***Admixture proportions*** | | | tdivadm<tdivebraf<br>tdivebram<tdivamaf<br>tdivebram<tdivebraf<br>tadmebro<tdiv1<br>tadmebro<tdiv2<br>tadmebro<tdiv1b<br>tadmebro<tdiv2b<br>tdiv1b>tdiv2b<br>tdiv1<tdiv2<br>tct2<tdivamaf2<br>tct2<tdivebram<br>tdivamaf2<tdivebraf<br>tct3<tct2 | | |
| ra | 0.01 | 0.99 | | | |
| ra2 | 0.01 | 0.99 | | | |
